# A Single Defined Sister Chromatid Fusion Destabilizes Cell Cycle through Micronuclei Formation

**DOI:** 10.1101/607341

**Authors:** Katsushi Kagaya, Naoto Noma, Io Yamamoto, Sanki Tashiro, Fuyuki Ishikawa, Makoto T Hayashi

**Affiliations:** The Hakubi Center for Advanced Research, Kyoto University, Yoshida-Konoe-cho, Sakyo-ku, Kyoto 606-8501, Japan; Seto Marine Biological Laboratory, Field Science, Education and Reseach Center, Kyoto University, Shirahama-cho, Nishimuro-gun, Wakayama 649-2211 Japan; Department of Gene Mechanisms, Graduate School of Biostudies Kyoto University, Yoshida-Konoe-cho, Sakyo-ku, Kyoto 606-8501, Japan

## Abstract

Chromosome fusion is deleterious among oncogenic chromosome rearrangements, and has been proposed to cause multiple tumor-driving abnormalities. Conventional methodologies, however, lack the strictness of the experimental controls on the fusion such as the exact timing, the number and the types of fusion in a given cell. Here, we developed a human cell-based sister chromatid fusion visualization system (FuVis), in which a single defined sister chromatid fusion is induced by CRISPR/Cas9 concomitantly with mCitrine expression. Fused chromosome developed numerical and structural abnormalities, including chromosome fragmentation, an indicative of eventual chromothripsis. Live cell imaging and hierarchical Bayesian modeling indicated that micronucleus (MN) is generated in the first few cell cycle, and that cells with MN tend to possess cell cycle abnormalities. These results demonstrate that, although most cells can tolerate a single fusion, even a single sister chromatid fusion destabilizes cell cycle through MN formation.

## Introduction

Chromosome abnormalities are at the core of tumorigeneis. Among oncogenic chromosomal rearrangements, chromosome fusion that gives rise to dicentric chromosome, is highly deleterious because of the generation of unresolved chromatin bridges following anaphase [1]. Chromosome ends are protected from activating DNA damage response (DDR) pathway by telomere that is composed of repetitive DNA sequence and specific DNA binding proteins [1]. In most human somatic cells, telomeric DNA gradually shorten in the absence of telomerase, an RNA-dependent DNA polymerase specific to telomeric sequence. Telomere shortening eventually induces DDR due to insufficient length of telomeric repeats for the end protection. In the absence of functional DDR pathway, cells keep dividing and enter a stage called telomere crisis [2]. During telomere crisis, chromosome ends become too short to inhibit non-homologous end joining (NHEJ) [3], which results in chromosome end-to-end fusion and consequently massive cell death [4] [5]. On the other hand, chromosome fusion during telomere crisis also induces chromosome abnormalities, which potentially generate mutations that increase cellular fitness [6]. Accumulation of such mutations, including telomerase promoter mutations, has been proposed to develop minor survivors, which eventually become tumors [1].

Previous studies led to the hypotheses that chromosome fusions cause multiple tumor-driving abnormalities including breakage-fusion-bridge (BFB) cycle [7] [8], binucleation [9], chromothripsis and kataegis [10], and mitotic arrest [4]. In these studies, the effects of chromosome fusions have been analyzed by artificial disruption of telomere binding proteins that protect chromosome ends from activating DDR. Among telomere binding complex called shelterin, TRF2 is central in telomere protection and targeted by various methods including dominant negative allele [11], shRNA-dependent knockdown [12] [13], and cre-loxP- and CRISPR/Cas9-mediated knockout [14] [4]. The fate of chromosome fusion has also been analyzed during telomere crisis induced by replicative telomere shortening in p53-compromised cells and mice that lack functional telomerase [15] [16] [17] [18]. During telomere crisis, cells often possess a few chromosome fusions, which is much milder than the phenotype induced by complete TRF2 disruption [14] [4]; while ongoing telomere shortening gives rise to continuous emergence of dicentric chromosomes [16]. Thus, in both experimental systems, multiple chromosome fusions are induced over time to different extent. Besides, there are at least three different types of chromosome end-to-end fusion induced in these systems. Inter-chromosomal fusion involves chromosome ends of two distinct chromosomes, while intra-chromosomal fusion occurs between both ends of the same chromosome, resulting in ring-shaped chromosome. The third is sister chromatid fusion that requires each end of sister chromatid pair after DNA replication. Among these, sister chromatid fusion has been implicated in the escape from telomere crisis by inducing appropriate genetic alterations [19]. Conventional methodologies failed to regulate the number and the types of chromosome fusion; it was very difficult to know the types and the number of fusion in a given cell without harvesting the cell, and the exact timing at which it happened. Recently developed method that uses sequence specific nucleases such as I-SceI and TALEN to induce double strand break (DSB) in subtelomere region, can potentially regulate the number of fusion as a consequence of abnormal repair between two distinct subtelomeric DSB [20] [21]. However, nuclease-mediated method still failed to regulate the types of fusion and the timing of its induction. Especially, it was very difficult to track some stochastic events happening within the cell lineage after a single defined chromosome fusion due to the lack of a method to visualize such fusion.

To circumvent these difficulties, we have developed a cell-based sister chromatid fusion visualization (FuVis) system, by which a single defined sister chromatid fusion can be artificially induced in a traceable manner. The FuVis system relies on CRISPR/Cas9-mediated targeting of an artificial cassette integrated in the X chromosome short arm (Xp) subtelomere; DSBs induced by the CRISPR/Cas9 generate a sporadic sister chromatid fusion concomitantly with mCitrine expression. We have successfully isolated two independent FuVis clones for Xp sister chromatid fusion (FuVis-XpSIS). PCR-based sequence analysis of the fusion junction suggested microhomology-mediated repair mechanism is involved in the sister chromatid fusion. Cytological analysis revealed that mCitrine positive FuVis-XpSIS cells indeed possess sister chromatid fusion and numerical and structural X chromosome abnormalities, including translocation, inter-chromosomal fusion and X chromosome fragmentation, which potentially gives rise to chromothripsis [22]. We also found that mCitrine positive FuVis-XpSIS cells did not develop tetraploidy, suggesting that a single sister chromatid fusion did not inhibit cytokinesis but was rather resolved or broken after cytokinesis. Live cell imaging and a lineage tracking of the mCitrine positive FuVis-XpSIS and FuVis-XpCTRL cells suggested that individual lineages in each clone possess various cellular abnormalities, including interphase delay, mitotic delay, micronucleus (MN) formation, multinuclei formation, cell death, cytokinesis failure and cell fusion. Together with a possible effect of the sister chromatid fusion in a given cell, we explicitly incorporated unknown lineage individualities to statistical models as hierarchical structure of the parameters. Not only the hierarchical models, we also built up alternative models to examine the validity of our consideration formulated as model structures. We assessed the appropriateness of the models in terms of the predictability by WAIC (Widely Applicable Information Criterion), which is a statistical measure applicable in the Bayesian inference [23] [24]. Our analysis indicates that development of MN depends on the sister chromatid fusion independently of the lineage individuality and other experimental variables and occurs in the first few cell cycle upon its formation. The analysis also indicates that cells with MN tend to delay interphase of the cell cycle and possess more abnormalities compared to their MN-negative sister lineages. These results illuminate that FuVis is a very powerful tool to follow the fate of a single defined DNA rearrangement. We propose that, although most cells can tolerate a single fusion, even a single sister chromatid fusion causes deleterious effect on cellular fitness through MN formation.

## Results

### Induction of chromosome fusion by CRISPR/Cas9

Targeting two distinct loci on chromosomes by CRISPR/Cas9 causes sporadic translocation [25]. We reasoned that targeting subtelomere sequence by CRISPR/Cas9 results in sporadic chromosome end-to-end fusion and/or sister chromatid fusion by erroneous repair (Supplementary figure 1A). Because human subtelomere loci contain multiple repetitive sequences [26], we picked up a target sequence, which is found in multiple different subtelomere regions by using online CRISPR design tool [27] and Cas-OFFinder [28]. The sequence was cloned into lentiviral CRISPR/Cas9 vector (CRISPR/Cas9-sgSubtel, Supplementary file 1). Normal human lung fibroblast IMR-90 cells expressing papilloma virus oncoprotein E6 and E7, which downregulate p53 and Rb, respectively, were infected with lentiviruses carrying CRISPR/Cas9-sgSubtel and selected with puromycin from 2 days post infection for 5 days. Metaphase spread followed by fluorescent in situ hybridization (FISH) with centromere and telomere probes revealed that targeting subtelomere sequences induced inter-chromosome and sister chromatid fusions (Supplementary figure 1B).

### Development of sister chromatid fusion visualization system

Successful induction of sister chromatid fusions by CRISPR/Cas9 prompted us to develop a cellular system, in which sister chromatid fusion can be detected by expression of a fluorescent protein only upon CRISPR/Cas9-induced sister chromatid fusion formation. To achieve this system, we designed DNA cassette that harbors mCitrine gene that is separated by splicing donor and acceptor (Figure 1A). The N-terminus is driven by the EF-1 promoter, while C-terminus is located upstream of an EF-1 promoter in an opposite direction. The downstream of the N-terminus harbors a spacer region, which contains multiple CRISPR/Cas9 target sequences that are not found in the human genomic sequence, and splicing acceptor- and self-cleaving peptide sequence (P2A)-tagged neomycin resistant (neoR) gene (Figure 1A). We reasoned that, upon integration of the cassette into a single subtelomere locus, targeting the spacer region between the N-terminus of mCitrine and the neoR gene by CRISPR/Cas9 induces a sporadic sister chromatid fusion, which results in the expression of full length mCitrine gene (Figure 1B). For this purpose, the entire cassette sequence was flanked by tandem cHS4 insulators, which suppress spreading of silent chromatin structure [29], and integrated into a telomere-adjoining subtelomere locus on the short arm of X chromosome in HCT116 cells by homology-mediated recombination (HR) upon CRISPR/Cas9 targeting (Figure 1A and supplementary file 1). We performed two independent HR-mediated integration of the cassette and isolated several clonal cell lines by G418-selection (24 and 48 clones in the first and second integration, respectively). The integration of the cassette into Xp subtelomere was analyzed by genomic PCR (Supplementary figure 1C, D, and data not shown). We obtained 2 positive (clone1-15, 21) and 9 positive (clone2-3, 6, 9, 13, 21, 33, 36, 38, 39) clones from the first and second integration, respectively (Supplementary file 2). We performed a second screening by using CRISPR/Cas9 that targets the spacer region (Figure 1B), which resulted in the expression of mCitrine at various levels in individual clones (Figure 1C and Supplementary figure 1E, F). The clone1-15 and the clone2-36 gave the highest mCitrine expression in each independent integration. We named the system as Fusion Visualization system for Xp SISter chromatid fusion (FuVis-XpSIS) and renamed the clone1-15 and clone2-36 as (FuVis-)XpSIS15 and (FuVis-)XpSIS36 cells, respectively (collectively called XpSIS). The integration of the cassette into the Xp subtelomere was further confirmed by FISH (Figure 1D and Supplementary figure 2A) and by Southern hybridization (Supplementary figure 1G, H). To confirm the induction of sister chromatid fusion by CRISPR/Cas9 targeting, mCitrine positive XpSIS15 cells expressing CRIPSPR/Cas9-sgFUSION11 (XpSIS15-sgFUSION11) were sorted and analyzed by FISH. We found that telomere-free Xp sister chromatid ends were connected and colocalized with sister cassette signal (Figure 1E and Supplementary figure 2B), which indicates sister chromatid fusion at the cassette locus.

**Figure 1.**
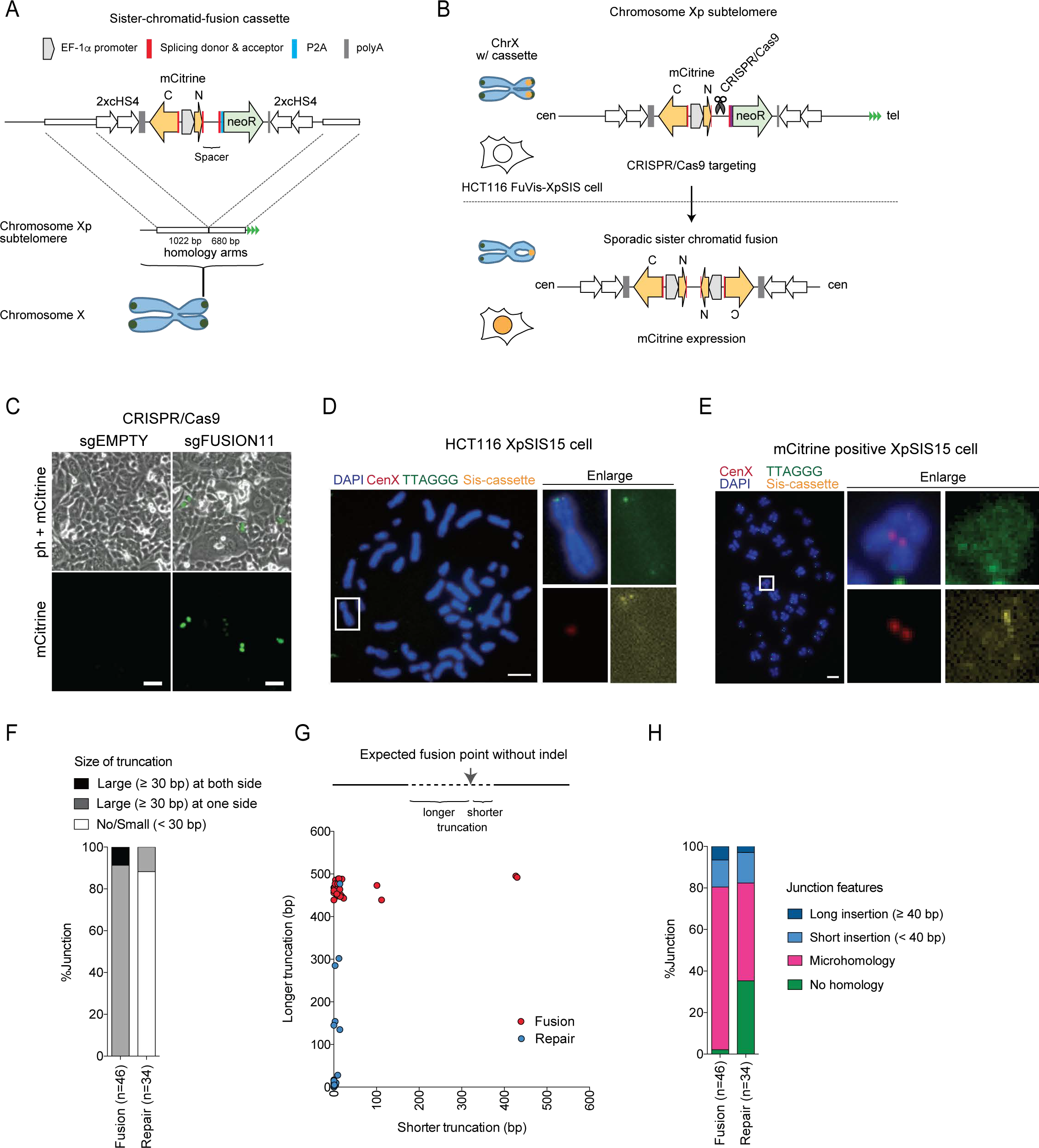
CRISPR/Cas9-mediated sister chromatid fusion visualization system. **A**, Schematic overview of the development of sister chromatid fusion visualization system. Sister chromatid fusion cassette harboring split mCitrine and the neoR gene flanked by tandem cHS4 insulator was integrated into Xp subtelomere in HCT116 cells by homology directed recombination. Clonal lines were selected by G418 and named as FuVis-XpSIS cell lines. **B**, Schematic representing CRISPR/Cas9-mediated induction of sister chromatid fusion and mCitrine expression. The spacer region between N-terminus of mCitrine and the neoR is targeted by CRISPR/Cas9, which results in sister chromatid fusion and full length mCitrine expression. **C**, Phase-contrast and fluorescence images of XpSIS36 cells harboring lentivirus-delivered CRISPR/Cas9-sgEMPTY or CRISPR/Cas9-sgFUSION11 at 10 days post infection. Scale bar, 50 *µ*m. **D, E**, FISH image of XpSIS15 (D) and XpSIS15-sgFUSION11 (E) showing X chromosome centromere (red), telomere (green), the sister cassette (yellow) and DAPI (blue). Scale bar, 10 *µ*m. mCitrine positive cells were sorted 8 days after CRISPR/Cas9-sgFUSION11 infection in (E). **F**, Percentage of truncation features at the fusion and repair junction in XpSIS15-sgFUSION11. The numbers of sequenced clones are indicated. **G**, Distribution of truncated length at the fusion and repair junction in (F). **H**, Percentage of the fusion and repair junction features in XpSIS15-sgFUSION11. Microhomology indicates the junction has 1-5 bp microhomology. Short and long insertion indicate insertion of other chromosomal sequence at the junction.

To evaluate the efficiency of mCitrine expression by targeting different sequences, XpSIS cells were separately infected by multiple lentiviruses containing different CRISPR/Cas9-sgFUSIONs, which target distinct loci in the spacer and the neoR gene (Supplementary file 1). We found that several CRISPR/Cas9-sgFUSIONs induced mCitrine expression and that the absolute level of mCitrine expression was higher in XpSIS36 than that of XpSIS15 across the entire target sites, while the relative expression of mCitrine among different target sites was similar between two clones (Supplementary figure 1I). Among these target sequences, sgFUSION11 was the most efficient inducer of mCitrine in both clones and chosen for the subsequent analysis unless otherwise indicated.

### Sister chromatid fusion junction analysis

To confirm that mCitrine positive XpSIS cells harbor covalently linked sister chromatid fusion, genomic DNA was extracted from mCitrine positive XpSIS15-sgFUSION11 cells and subjected to PCR amplifying expected fusion junction (Supplementary figure 2C). PCR product specific to mCitrine positive XpSIS15 was shorter than expected size (Supplementary figure 2D and data not shown), which was then cloned for subsequent sequencing analysis. Cloned junction sequences were successfully aligned to the expected junction, although at least one side of the junction was truncated longer than 30 bp in all clones (Figure 1F, G; Fusion), indicating large resection upon sister chromatid fusion. To address if a repair process in Xp subtelomere region always coincides with large truncation, expected repair product upon CRISPR/Cas9-targeting was also amplified and cloned using genomic DNA from entire population of XpSIS15-sgFUSION11 cells (Supplementary figure 2E). Alignment of these clones showed that about 90% of clones do not possess large truncation (Figure 1F, G; Repair), suggesting that large resection is specific to sister chromatid fusion process. The sister chromatid fusion junction preferred microhomology while repair products showed more junctions without any homology (Figure 1H). Fusion junction in XpSIS15-sgFUSION4 and XpSIS36-sgFUSION11 also possessed similar features (Supplementary figure 2F-I), excluding the possibility that the observed features are specific to a certain target sequence and/or XpSIS clone. The preference of large truncation and microhomology at the fusion junction suggests that sister chromatid fusion is processed by microhomology-mediated end joining (MMEJ), which is active in late S/G2 phase when sister chromatids are present [30]. This profile of the fusion junction is consistent with naturally occurred and TALEN-induced chromosome end-to-end fusions [31] [32] [33] [21].

### Construction of the control system

Since the XpSIS relies on targeting of the subtelomeric cassette by CRISPR/Cas9 and the resulting expression of mCitrine, we designed a control system, in which CRISPR/Cas9 targeting of Xp subtelomere induces mCitrine expression without sister chromatid fusion (Figure 2A, B). For this purpose, the splicing-acceptor-tagged neoR gene and two tandem polyA sequences were inserted between the N-terminus and the C-terminus of mCitrine in the same direction (Figure 2A). Both upstream and downstream of the neoR gene contain the same sgFUSION11 target sequences so that CRISPR/Cas9 targeting is expected to result in a sporadic deletion of the neoR gene and mCitrine expression (Figure 2B). Among 48 G418-resistant clones obtained upon integration of the control cassette into Xp subtelomere, 3 clones (named as FuVis-XpCTRL16, 33 and 48) showed genomic PCR product suggesting integration (Supplementary figure 3A, B and data not shown)(Supplementary file 2). XpCTRL48 gave a PCR product longer than expected, which was confirmed to be caused by a duplication of telomere-distal homology arm sequence, while XpCTRL16 and XpCTRL33 showed higher background mCtirine expression (supplementary file 2 and data not shown); therefore we chose XpCTRL48 cells for subsequent analysis. The integration was further confirmed by Southern hybridization (Supplementary figure 3C, D) and FISH (Figure 2C and Supplementary figure 3E). XpCTRL48 cells also possess background level of mCitrine expression by FACS analysis (Figure 2E). However, introduction of CRISPR/Cas9-sgFUSION11 into XpCTRL48 induced robust expression of mCitrine (Figure 2D,E). We performed sequencing analysis of repair junctions using genomic DNA from whole population of XpCTRL48-sgFUSION11 (Figure 2F). The PCR generated a long product which corresponds to the original sequence and faint short product which was expected to contain truncated repair product (Figure 2G); the latter was cloned for the sequencing analysis. The sequenced clones possessed two types of junction: those that completely lost sequences between CRISPR/Cas9 targets (type I) and that partially lost the sequences (type II)(Figure 2H). The type II suggests that each CRISPR/Cas9 target site was independently repaired, while the type I contains either product of sequential repair of each site or direct repair between the two target sites (Figure 2H). Both types of junction possessed a sign of MMEJ, which was indicated by extensive truncation, microhomology and insertions at the junction (Figure 2I, J) [34] [19]. These results suggest that both XpSIS and XpCTRL (collectively called FuVis-Xps) rely on MMEJ for the targeted genome rearrangements required for mCitrine expression.

**Figure 2.**
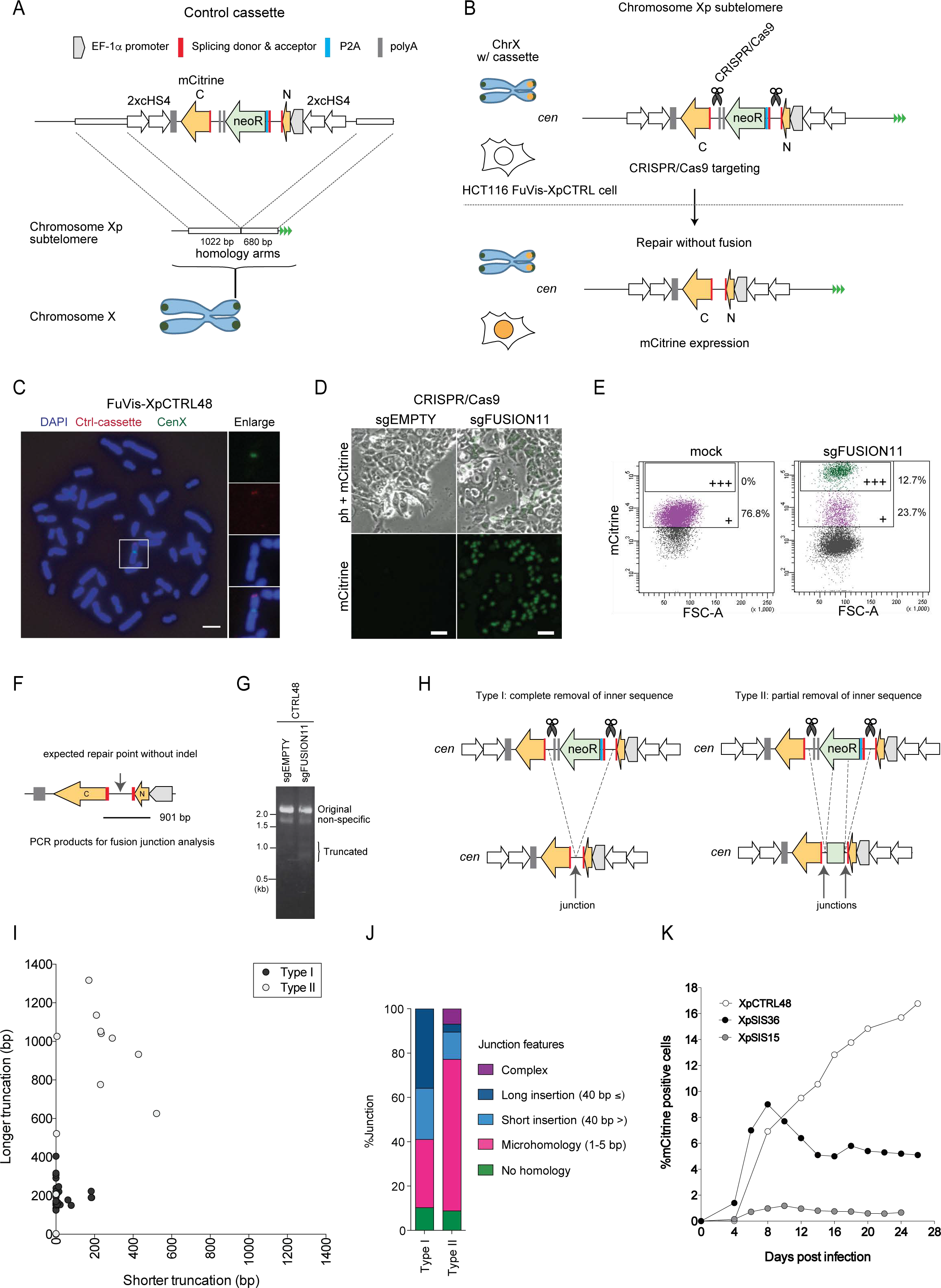
CRISPR/Cas9-mediated mCitrine expression system as a control. **A**, Schematic overview of mCitrine expression control system. A control cassette harboring the neoR gene flanked by the split mCitrine and tandem cHS4 insulators was integrated into Xp subtelomere of HCT116 by homology directed recombination. **B**, Schematic representing CRISPR/Cas9-mediated removal of the neoR gene and mCitrine expression. Spacer regions flanking the neoR gene harbor the same CRISPR/Cas9 target sequences. Sporadic sister chromatid fusion can also occur but does not induce mCitrine expression. **C**, FISH images of XpCTRL48 cells showing X chromosome centromere (green), the control cassette (red) and DAPI (blue). Scale bar, 10 *µ*m. **D**, Phase-contrast and mCitrine fluorescence images of XpCTRL48-sgEMTPY or -sgFUSION11 cells at 10 days post infection. Scale bar, 50 *µ*m. **E**, Flow cytometory analysis of XpCTRL48 cells in (D). Symbols “+” and “+++” indicate background level of mCitrine expression and fully expressed mCitrine following CRISPR/Cas9-induced removal of neoR gene, respectively. **F**, Schematic representing a PCR product used for repair junction analysis in XpCTRL48-sgFUSION11. **G**, Results of PCR shown in (F) using indicated genomic DNA. **H**, Schematics for two types of repair junction. The type I possesses complete removal of the neoR sequence between Cas9-target sites; the type II shows partial removal and suggests sequential repair of each junction. **I**, Distribution of truncated length across repair junctions in the type I and II. **J**, Percentage of junction features in the type I and II as shown is Figure 1H. Complex indicates junction that has multiple independent insertions of original cassette sequence. **K**, Time course of mCitrine expression in Xps-sgFUSION11. The result was reproducible in two independent experiments.

### Kinetics of mCitrine expression upon CRISPR/Cas9-targeting

To analyze the kinetics of mCitrine expression following sister chromatid fusion, Xps-sgFUSION11 were cultured for more than three weeks post infection. Flow cytometry analysis revealed that percentage of mCitrine positive cells gradually increased in XpCTRL48 during entire culture period, while it peaked at 8 and 10 days post infection in XpSIS36 and XpSIS15, respectively, and gradually decreased until it reached a plateau around 14 days post infection (Figure 2K). This kinetics is consistent with an assumption that, upon sister chromatid fusion, a single mCitrine gene is generated in G2 phase, which can be propagated to either one of two daughter cells following the first mitosis, while repair in XpCTRL resulted in two mCitrine genes in G2 phase (Supplementary figure 3F, G). To address this assumption, mCitrine-positive Xps-sgFUSION11 cells at day 8 post infection were sorted, cultured for another 13 days and analyzed by FACS again (Supplementary figure 3H). Strikingly, 94% of XpCTRL48-sgFUSION11 cells were still mCitrine positive, while only 42% of XpSIS-sgFUSION11 cells were mCitrine positive (Supplementary figure 3I, J), indicating that mCitrine gene was not propagated to both daughter cells in XpSIS-sgFUSION11 cells.

### Numerical and structural abnormality of X chromosome induced by sister chromatid fusion

To measure X chromosome abnormality upon sister chromatid fusion, dual-colored FISH was performed on metaphase spreads using DNA probes spanning the entire X chromosome (red) and the X centromere (green). mCitrine positive XpSIS36-sgFUSION11 and XpCTRL48-sgFUSION11 cells were sorted at day 8 and 14 post infection (D8 and D14) and analyzed by metaphase spread (Figure 3A). Besides, mCitrine positive cells at D8 were sorted and re-cultured for another 6 days (D8-14) to follow the long-term fate of X chromosome sister chromatid fusion (Figure 3A). X chromosome remained mostly intact in uninfected XpSIS36 and XpCTRL48 cells, and mCitrine positive XpCTRL48-sgFUSION11 cells (Figure 3B). In contrast, mCitrine positive XpSIS36-sgFUSION11 cells possessed both numerical and structural X chromosome abnormalities (Figure 3C,D). To assess numerical abnormality, total number of chromosomes, a percentage of near tetraploid cells and a ratio of X chromosome per total chromosomes in each cell were plotted. Total chromosome number and the percentage of near tetraploid cells did not change in all conditions (Figure 3E and Supplementary figure 4A), suggesting that a single sister chromatid fusion did not cause cytokinesis failure and binucleation. XpSIS36 cells treated with a myosin II inhibitor Blebbistatin for 48 hr showed accumulation of near tetraploid and octaploid cells, excluding the possibility that tetraploid cells could not enter mitosis (Figure 3E and Supplementary figure 4A). On the other hand, the ratio of X chromosome per total chromosomes became variable at D8 in XpSIS36-sgFUSION11 cells compared to other conditions (Figure 3F), indicating that X chromosome was mis-segregated during mitosis. This variability became less obvious at D14 and after re-culturing (D8-14), suggesting that cells with X chromosome aneuploidy had been removed from the population. Figure 3G represents percentage of X chromosome structural abnormalities, which include translocation, inter-chromosome fusion with non-X or X chromosome, and fragmented chromosome, and became prominent only in XpSIS36-sgFUSION11 cells (Figure 3D, G and Supplementary figure 4B). After re-culturing, 38.9% and 51.1% of cells possessed translocation and seemingly normal X chromosome, respectively. These results suggest that, in most cells, sister chromatid fusion has been broken after first mitosis and stabilized by either translocation of other chromosome portion that contains functional telomere or de novo telomere addition, a phenomenon called telomere healing [35]. Inter-chromosome fusion with non-X was observed at all time points (D8, 10%; D14, 5.6%; D8-14, 5.5%), suggesting ongoing BFB cycle in small population of the cells. Fusion between two X chromosome was rare and observed even in the untreated cells (mock, 1.1%; D8, 1.1%; D14, 0%; D8-14, 3.3%). However, while it was observed in near-tetraploid cells in mock control, near-diploid cells showed the phenotype in XpSIS36-sgFUSION11 condition and such X-X fusion contributed to X chromosome aneuploidy, which implies different mechanism of origin. The fragmentation was also rare (D8, 5.5%; D14 and D8-14, 1.1%) but specific to XpSIS36-sgFUSION11, suggesting that the phenotype was induced by the X sister chromatid fusion.

**Figure 3.**
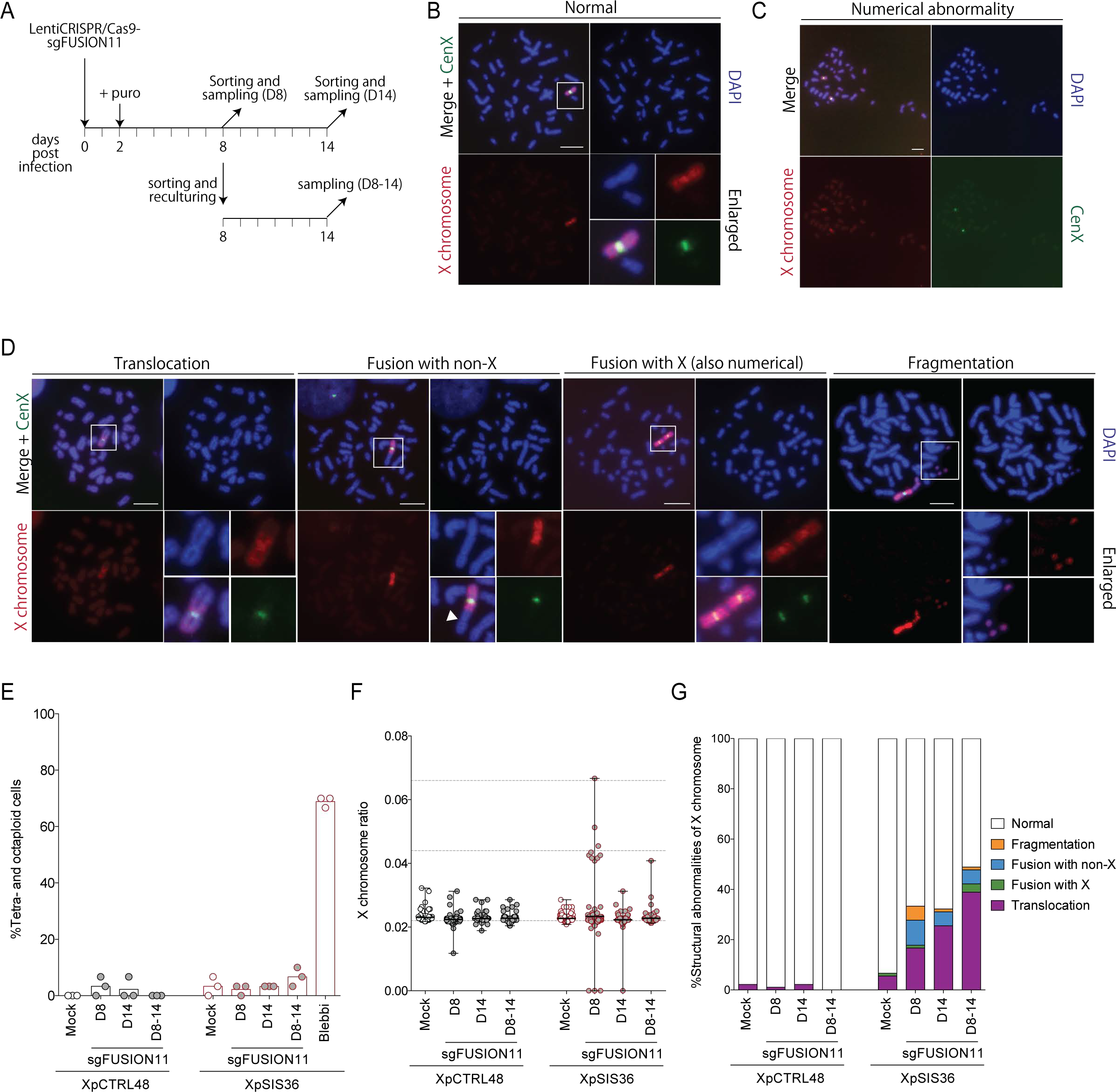
A single sister chromatid fusion causes numerical and structural abnormalities on X chromosome. **A**, Schematic of FISH analysis. XpCTRL48 and XpSIS36 cells infected with CRISPR/Cas9-sgFUSION11 lentivirus were selected with puromycin. mCitrine-positive cells were sorted at 8 and 14 days post infection for FISH analysis (D8 and D14). Besides, mCitrine positive cells sorted at D8 were re-cultured for 6 days and harvested (D8-14). **B-D** Representative FISH images of XpSIS36-sgFUSION11 cells showing whole X chromosome (red), X chromosome centromere (green) and DAPI (blue). Scale bar, 10 *µ*m. Shown are normal metaphase spread of mock treated cells (B), numerical abnormality (C), and structural abnormalities (D). White arrowhead in fusion with non-X indicates a narrow region on non-X chromosome, an indicative of centromere locus. **E**, Percentage of near-tetraploid and near-octaploid mitosis in indicated cells (n=3). Blebbi, 100 *µ*M blebbistatin for 48 hr. **F**, Ratio of X chromosome per total chromosomes in indicated cells. Results of 90 spreads from three independent experiments are shown with mean and range. **G**, Ratio of structural abnormalities in indicated cells. Thirty metaphase spreads were analyzed per experiment. Mean of three independent experiments are shown.

### lineage analysis of mCitrine positive cells by live cell imaging

To further dissect the cellular events following sister chromatid fusion, we performed a lineage analysis by live cell imaging of FuVis-Xps. For each cell line, we generated control cells expressing full-length mCitrine introduced by retrovirus infection to characterize cell cycle properties of each cell line under mCitrine expression without CRISPR/Cas9 targeting (Figure 4A). These cell lines are named as XpCTRL48-mCit, XpSIS15-mCit and XpSIS36-mCit (collectively Xps-mCit). In Xps-sgFUSION11, we observed two types of mCitrine positive cell: those that became mCitrine positive during the course of the movies; and those that were already mCitrine positive at the beginning of the movies (Figure 4B). The former represents the first few rounds of cell cycle after CRISPR/Cas9-targeting and repair (XpCTRL48-sgFUSION11) or sister chromatid fusion formation (XpSIS-sgFUSION11), while the latter represents (N+n)th rounds of the cell cycle after the repair or the fusion events (Figure 4A,B).

**Figure 4.**
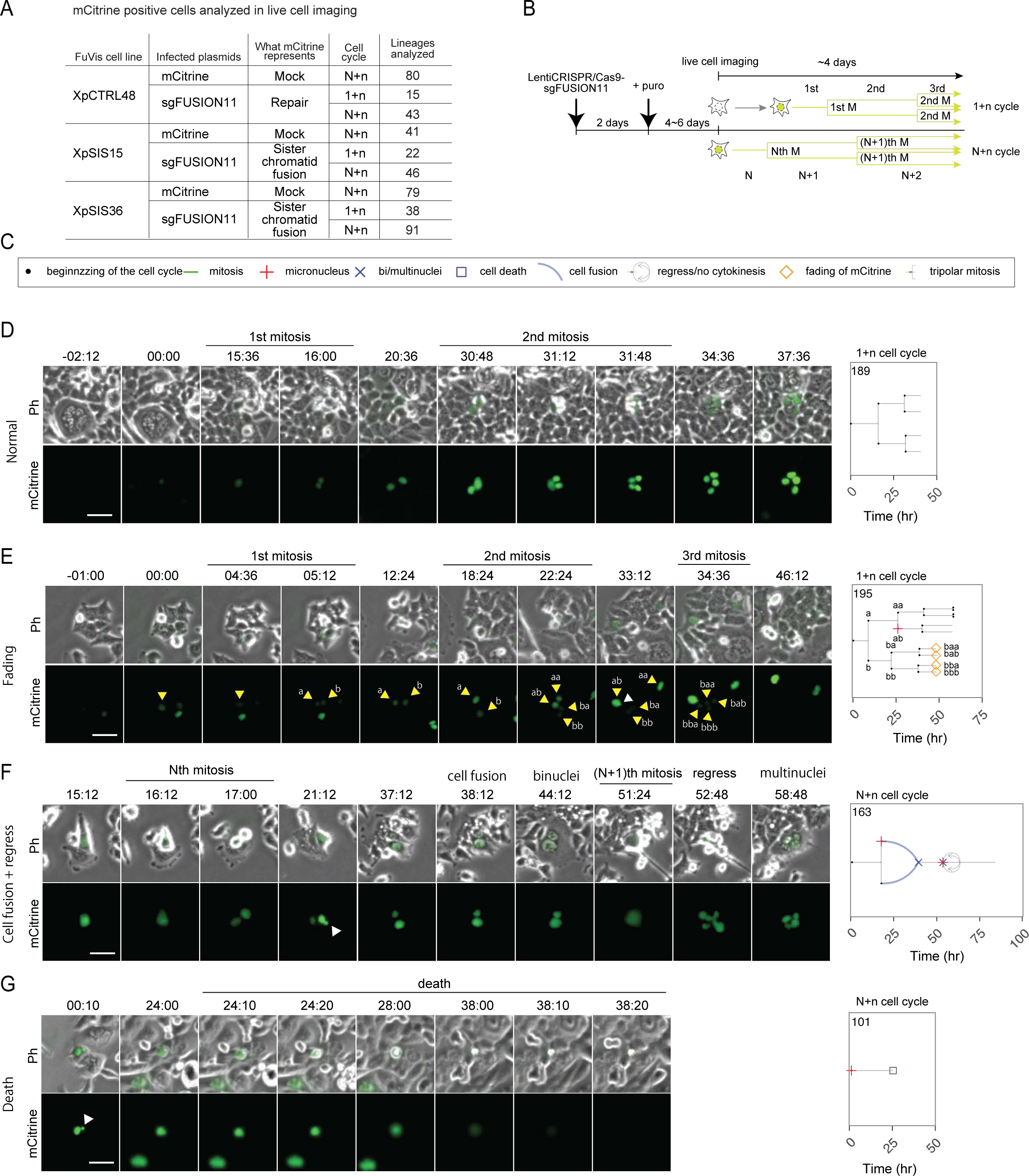
Live cell analysis of the fate of individual FuVis-Xps cells expressing mCitrine. **A**, A table representing FuVis cells analyzed in live cell imaging. Infected plasmids were either retroviral pMX-mCitrine or lentiviral CRISPR/Cas9-sgFUSION11. Cell cycle indicates the number of cell cycle after mCitrine expression as shown in (B). The numbers of lineages analyzed in each condition are indicated. **B**, Schematic of live cell imaging. Typical live cell imaging was started after 6 to 8 days post infection, when puromycin selection had been completed. Read the main text for the detail. **C**, Symbols representing cell cycle progression and cellular abnormalities in lineage trees. **D-G**, Representative live cell images of the fate of mCitrine positive XpSIS36-sgFUSION11 cells (left) and corresponding lineage trees (right). Shown images are normal cell division during 1+n cell cycle (D), fading of mCitrine in one of two sister cell lineages during 1+n cell cycle (E), sister cell fusion followed by cytokinesis failure during N+n cell cycle (F), and cell death in Nth cell cycle (G). Cells in the same lineage are indicated by yellow arrowhead with alphabets (a, b, aa, ab, ba, bb, baa, bab, bba and bbb) denoting a lineage order in (E). White arrowhead indicates MN. Scale bar, 50 *µ*m. In the lineage trees, the time points of cell division are marked by bifurcation with green bars representing mitotic duration. When an individual cell shows a sign of MN or bi/multinuclei formation, the beginning of the cell cycle is marked by the representing symbols. When a given cell show a sign of cell death or fading of mCitrine, the time point of the events is marked by the representing symbols. Blue lines represent cell fusion events, which can occur both in inter- and intra-lineage manner.

We have manually followed lineages of each mCitrine positive cell and analyzed the following characteristics: interphase duration, mitotic duration, a fading of mCitrine and cellular abnormalities. The observed cellular abnormalities include cell death, MN formation, bi/multinuclei formation, tripolar mitosis, cytokinesis failure/furrow regression, and cell fusion. The number of lineages analyzed are summarized in Figure 4A. Cell cycle progression of individual lineages of mCitrine positive cells were visualized as lineage trees with distinct symbols representing the cell cycle features (Figure 4C). For example, figure 4D shows live cell image of the first few cell cycle of a normal cell division in XpSIS36-sgFUSION11. The corresponding lineage tree is shown on the right, which represents interphase duration, mitotic duration and cell division by a gray bar, a green bar, and a bifurcation, respectively. Typical examples of a fading of mCitrine in 1+n cell cycle (Figure 4E), daughter cell fusion followed by regression and multinucleation in N+n cell cycle (Figure 4F) and cell death in N+n cell cycle (Figure 4G) are also represented. All lineage trees analyzed in this study are shown in Supplementary figures 5-13.

### Modeling of the fate of a single sister chromatid fusion

To assess the consequence of a single sister chromatid fusion, percentages of lineages that show cell cycle and morphological abnormalities were calculated from the lineage trees and shown as a heat map (Figure 5A and Supplementary figure 14A). No mitosis indicates a lineage that did not enter mitosis during the course of the movie without any sign of cell death, fading of mCitrine, and/or cell fusion; mitotic delay was defined as mitosis that was longer than 2 hours. The fading of mCitrine was observed only in XpSIS-sgFUSION11, further confirming that this is the consequence of sister chromatid fusion (Supplementary figure 3F). Among all abnormalities assessed, lineages that are positive for MN formation became prominent in XpSIS-sgFUSION11 compared to other conditions (Figure 5A,B and Supplementary figure 14A-C).

**Figure 5.**
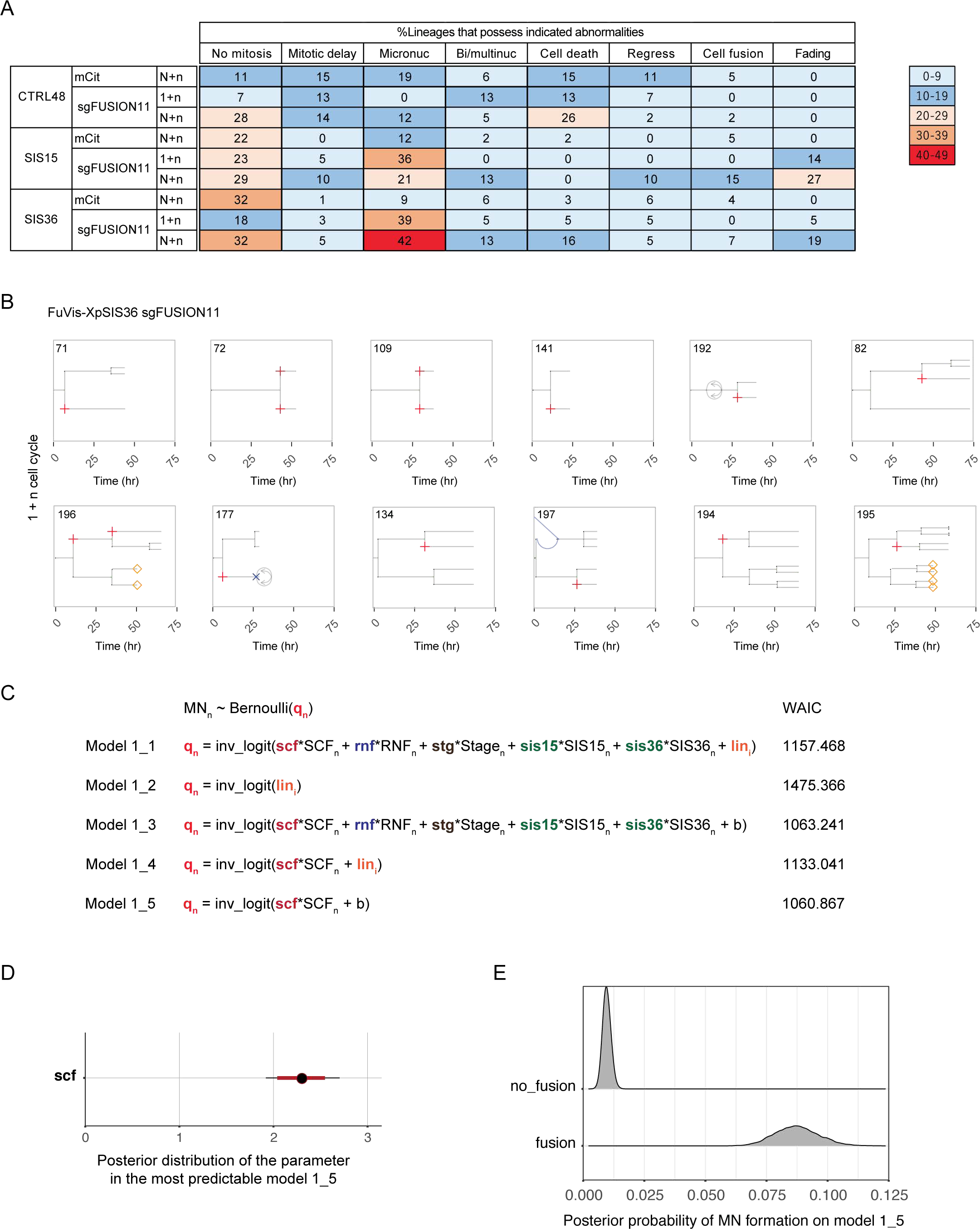
A single sister chromatid fusion generates micronucleus. **A**, A heat map representing the percentages of lineages that possess indicated cell cycle and morphological abnormalities in each condition. No mitosis indicates a lineage that did not enter mitosis during the course of the movie. Mitotic delay represents a lineage that possesses at least one mitosis longer than 2 hours. **B**, Lineages that show MN formation during 1+n cell cycle in XpSIS36-sgFUSION11. The numbers above the trees represent lineage ID. **C**, Five model structures constructed to explain the probability of micronucleus (MN) formation (*q*_*n*_), which is subjected to Bernoulli distribution (top). The *q*_*n*_ is modeled by the following function and parameters: *inv logit*, inverse logit function; *scf*, the coefficient of sister chromatid fusion; *rnf*, the coefficient of repair (no fusion); *stg*, the coefficient of cell cycle stage after mCitrine expression (1+n or N+n); *sis*15 and *sis*36, the coefficient of cell line (SIS15 or SIS36 compared to CTRL48); *lin_i_, i* = 1, …, *N*_*lineages*_, intercepts representing individuality (unknown cellular characteristics shared in each lineage); and *b*, a bias parameter. Large capitals indicate variables (dummy variables) obtained from data (0 or 1 in supplementary file 8). The models implement all explanatory variables and individuality (1 1), only individuality (1 2), all explanatory variables without individuality (1 3), only *scf* and individuality (1 4), and only *scf* (1 5). The calculated WAIC values for each model are shown on right. **D**, The posterior distribution of the parameter *scf* inferred from the most predictable model (model 1_5) with median (black circle), and 50% (red bar) and 95% (black bar) credible intervals. **E**, Distribution of posterior predicted probabilities of MN formation calculated by using the parameters inferred in the model 1_5. No fusion contains the following conditions: C48-mCit (1 + *n* and *N* + *n*), C48-sgFUSION11 (1 + *n* and *N* + *n*), SIS15-mCit (1 + *n* and *N* + *n*), and SIS36-mCit (1 + *n* and *N* + *n*). Fusion contains the following conditions: SIS15-sgFUSION11 (1 + *n* and *N* + *n*) and SIS36-sgFUSION11 (1 + *n* and *N* + *n*).

To statistically infer the impact of sister chromatid fusion on MN formation in individual cells, we performed the logistic regression, which is a kind of generalized linear models (GLMs) [36]. We modeled the probability of MN formation (responsive variable) as a function of the linear predictor including the following explanatory variables and their coeffcient parameters: SCF, sister chromatid fusion (XpSIS-sgFUSION11); RNF, repair/no fusion (XpCTRL48-sgFUSION11); STG, cell cycle stages after mCitrine expression (1+n or N+n); and SIS15 and SIS36, cell lines (XpSIS15 or XpSIS36 compared to XpCTRL48)(Figure 5C and supplementary file 3).

We also realized that, even in Xps-mCit, individual lineages in each cell line can possess distinct and diverse cell cycle characteristics (Supplementary figures 5-13), which potentially lead to over- or underestimation of the effect of the explanatory variables. We therefore examined the model including the individuality of cell lineages as hierarchical structure into the parameters (lin; unknown individuality of each lineage) (Figure 5D, model 1 1). Besides this model, models only with individuality (model 1 2), with all parameters except for individuality (model 1 3), only with SCF and individuality (model 1 4), and only with SCF (model 1 5) were also constructed and applied to the data obtained from movie analysis to estimate the posterior distribution of the parameters (supplementary file 3). We constructed the posterior predictive distributions and inferred the prediction errors of the models by WAIC. WAIC is a statistical measure based on a rigorous mathematical theory to estimate the generalization loss of models even if the posterior distribution is far from any normal distribution [23] [24]. Generally, the posterior distributions of our models with hierarchical structure do not resemble normal distribution. Therefore, we used this information criterion. Smaller the WAIC value is, more predictable the model is when a new data is generated.

The assessment of the models by WAIC demonstrated that the most predictable model is the model 1 5 (Figure 5C and supplementary file 3), indicating that implementation of neither lineage individuality nor linear predictors other than SCF improved the predictability of the model. The estimated posterior distribution of the parameter *scf* in the model 1 5 indicates that the median and 2.5 percentile of it are 2.29 and 1.92, respectively (Figure 5D). Given that if a parameter value is close to 0, the corresponding explanatory variable does not have an effect on a responsive variable, this result indicates that SCF has a positive effect on MN formation. Figure 5E shows distribution of posterior probability of MN calculated by the inferred value of the parameter *scf* in the model 1 5. The average posterior probabilities in the absence and the presence of sister chromatid fusion are 0.0096 and 0.087, respectively, suggesting that sister chromatid fusion increases the probability of MN by 9 times on average. We also estimated the distribution of the parameters in the second predictable model 1 3, which indicates the positive effect of sister chromatid fusion, while repair, cell cycle stage, and cell line were estimated to have relatively minor, if any, effects on the probability of MN formation (Supplementary figure 14D). Average posterior predicted probabilities of MN formation in the model 1 3 also indicated that sister chromatid fusion increases the expected probability of MN (Supplementary figure 14E). Thus, the predicted probabilities of MN in both models 1 3 and 1 5 indicate that MN is induced in the first few cell cycle upon a single sister chromatid fusion.

We asked if MN formation had any negative effect in the descending lineage. To address this question, two matched sister cells, in which one of them possesses MN, were picked up from all lineage trees (for example, Figure 5B and Supplementary figure 14C: all lineages except for 72 and 109), and their fates were compared. Our statistical inference indicated that cells with MN possess higher probability of subsequent abnormalities (0.271), including less frequency of mitosis, regress, cell fusion, cell death, and mitotic delay, than MN-negative sister lineages (0.063)(Figure 6A).

**Figure 6.**
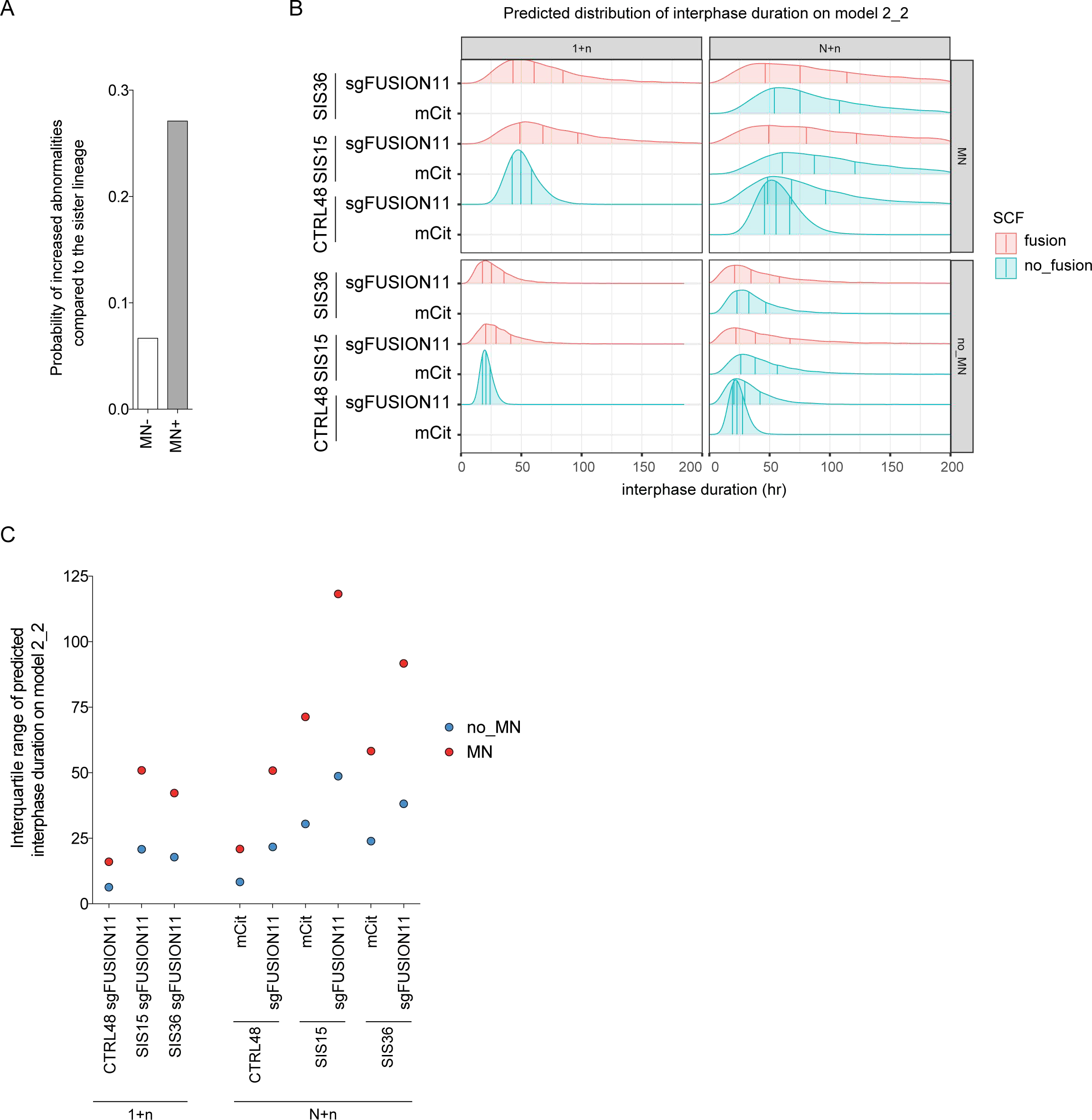
Micronucleus induced by a single sister chromatid fusion destabilizes the cell cycle. **A**, Bar graph shows the inferred probability of increased abnormalities compared to the sister lineage. Forty-eight matched sister pairs, one of which possesses MN, were selected from all lineage trees, and cell cycle abnormalities were compared between the sisters. The abnormalities include less mitosis in descending lineage, mitotic delay, cell death, regress, and cell fusion. **B**, Predicted distribution of interphase duration on the model 2_2. The values of parameters and sigma inferred from the most predictable model 2_2 were used to estimate distribution of interphase duration in indicated conditions. Conditions of the cell cycle stage (1+n or N+n) and MN (MN or no MN) are separately shown. Cells with sister chromatid fusion are represented in red color. The vertical bars indicate 25, 50, and 75 percentiles (from left to right). **C**, Dot plots representing interquartile range (IQR) of predicted interphase duration on the most predictable model 2_2. IQR was calculated by subtracting 25 percentile from 75 percentile in each condition in (B).

We presumed that cell cycle abnormalities would be detected as increased variability of cell cycle duration, because cells with the abnormalities tended to indicate an interphase delay. To assess the effect of MN formation on cell cycle duration, we made models assuming that the interphase duration is subjected to log-normal distribution, where the log of median (mu) is a function of the presence of MN and other variables (Supplementary figure 15A and supplementary file 3). Because both MN and interphase duration can be affected by the other variables (SCF, RNF, SIS15, SIS36, and STG), we assumed that the coefficient parameters (*scf*, *rnf*, *sis*15, *sis*36, and *stg*) are confounding factors (Supplementary figure 15B). The other parameter sigma was assumed to differ among each experimental condition. Models 2_1 and 2_2 implemented the presence of MN and all other variables with or without lineage individuality, respectively; a model 2_3 removed MN from the model 2_2; a model 2_4 implemented only MN, SCF and lineage individuality; a model 2_5 implemented only lineage individuality; and a model 2_6 implemented only MN and SCF (Supplementary figure 15A). The best performed model among the six models in terms of WAIC was the model 2_2, which implemented all the confounding factors. Therefore, our assumption about confounding factors to infer the causality between MN and interphase duration is in line with the model selection in terms of WAIC (Supplementary figure 15A,B and supplementary file 3).

The posterior distribution of the coefficient parameter of MN (micro) in the model 2_2 shows that the median and 2.5 percentile of it are 0.88 and 0.73, respectively (Supplementary figure 15C). Thus, it distributes far from zero on the model and the presence of MN would prolong interphase duration 2.4 (= *exp*(0.88)) times longer than its absence when the other explanatory variables are fixed. Predicted distribution of interphase duration in the presence and absence of MN in each experimental condition were calculated by using all posterior parameter distributions (Figure 6B and supplementary file 3). The shapes of distribution, shown with 25, 50, and 75 percentile bars, suggest that the presence of MN delays and broadens the distribution of interphase duration. To assess the variability of interphase duration, we plotted interquartile range (IQR) of each distribution (Figure 6C). Comparison of the IQR indicates that the presence of MN increases the variability of interphase duration regardless of the background conditions, and that SIS15-sgFUSION11 N+n and SIS36-sgFUSION11 N+n show the first and second highest IQR, respectively. These results and analyses support the notion that cells with MN induced by sister chromatid fusion possess the strongest destabilization of the cell cycle.

## Discussion

Chromosome fusions have been linked to multiple different abnormalities including anaphase bridge, cytokinesis failure, binucleation, chromothripsis, kataegis, mitotic arrest and cell death. These abnormalities are probably the consequences of multiple different types of chromosome end-to-end fusion through multiple rounds of cell cycle because conventional methods could not regulate the severity, the types and the exact timing of fusion. Thus, it was not possible to address the consequence of a single defined chromosome fusion especially in the first few cell cycle upon induction. Here, we have developed the FuVis-XpSIS system, a very powerful tool that allows visualization of a cell that possesses a single sister chromatid fusion upon its induction (Figure 1). By using FuVis-Xps, we have shown that a single sister chromatid fusion causes MN formation in the first few cell cycle and multiple chromosome abnormalities including inter-chromosome fusion and chromosome fragmentation (Figure 3, 5).

### Construction and assessment of FuVis

We have successfully isolated two FuVis-XpSIS and one FuVis-XpCTRL clones. Both XpSIS and XpCTRL rely on CRISPR/Cas9-mediated digestion of the spacer region to induce genomic rearrangements, which result in the full length mCitrine expression through MMEJ-mediated repair pathway (Figure 1F-H, Figure 2I,J and Supplementary figure 2F-I). However, only XpSIS, but not XpCTRL, requires fusion between two sister chromatids to acquire such rearrangement. Consequently, mCitrine expression in XpSIS cells is tightly linked to sister chromatid fusion formation. The generation of sister chromatid fusion upon CRISPR/Cas9-targeting in XpSIS was confirmed by multiple different observations including direct visualization of sister chromatid fusion by metaphase FISH (Figure 1E and Supplementary figure 2B), fusion junction sequences (Figure 1F-H and Supplementary figure 2F-I), fading of mCitrine expression (Figure 2K, Figure 4E and Figure 5A) and X chromosome abnormalities (Figure 3). We assume that the percentages of cells that acquired expected rearrangement in XpSIS and XpCTRL were underestimated because extensive truncation at the fusion junction would result in sister chromatid fusion and repair without mCitrine expression. Indeed, background expression of mCitrine in XpCTRL48 cells was diminished by CRISPR/Cas9-sgFUSION11 in some cells (Figure 2E), suggesting a partial loss of mCitrine gene upon repair process in this population. Such potential underestimation, however, does not affect the interpretation of the results since we could focus our analysis exclusively on mCitrine positive cells, which are associated with desired sister chromatid fusion and repair products in XpSIS and XpCTRL, respectively.

### Implication of the statistical modeling

We have applied the GLMs to the live cell imaging data to implement multiple different experimental variables in the statistical models. The hierarchical Bayesian models were also constructed to implement unknown lineage individuality in the statistical models. The lineage individuality assumes that, if a given cell possesses certain abnormality, ascending and descending cells in the same lineage tend to show the same abnormality, which is a reasonable assumption but often ignored in other statistical analysis. This assumption is important to avoid over- or underestimation of the effect of variables of interest when experimental data have clustered structure [37]. However, comparison of the model 1_1 and 1_3, the model 1_4 and 1_5, the model 2_1 and 2_2, and the model 2_4 and 2_6 in terms of WAIC demonstrated that the implementation of the lineage individuality did not improve the predictability of the models. This result suggests that individual lineages do not have tendency to possess similar abnormalities, although it is also possible that the live cell data do not have enough lineages that possess multiple cell cycle with abnormalities to assess lineage individuality in our model (Supplementary figures 5-13). We have therefore focused on the most predictable model without individuality in both models 1 and 2, which infer the effect of experimental variables in the sense of minimizing prediction error. The most predictable models indicate that sister chromatid fusion, but not other experimental variables, increases the probability of MN (Figure 5C-E), and that MN delays and destabilizes interphase duration (Figure 6B,C and Supplementary figure 15). The average predicted probability of MN formation on the model 1_5 increased by 9.1 times (from 0.0096 and 0.087) (Figure 5E). Thus, the statistical inferences from the best model can be interpreted that a single sister chromatid fusion increase the probability of MN formation by about 9 times regardless of the cellular backgrounds in our experimental setting.

In the model 2, we could also assess the effect of MN on the stability of the cell cycle. We have used IQR of interphase duration as a measure of cell cycle stability because when a cell possesses a certain abnormality, the cell is assumed to delay or even halt the cell cycle, which broadens the distribution of interphase duration. The comparison of IQR of predicted interphase duration on the most predictable model 2_2 suggests that MN destabilizes the cell cycle especially in XpSIS-sgFUSION11 N+n conditions (Figure 6C). Because the IQRs in XpSIS-sgFUSION11 1+n conditions are relatively low, we assume that the negative effect of MN induced by sister chromatid fusion accumulates during cell cycle progression. It is also possible that sister chromatid fusion possesses negative effect on cell cycle independently of MN formation during long-term culturing. The statistical analysis thus indicates that a single sister chromatid fusion, but not repair or other cellular characteristics, destabilizes the cell cycle through cumulative effects of MN formation and potentially other cellular abnormalities including BFB cycle, missegregation, and/or chromosome fragmentation.

### The fate of a single sister chromatid fusion on X chromosome

The results obtained from live cell imaging and X chromosome FISH in mCitrine positive FuVis-Xps allow us to model the fate of a single sister chromatid fusion in HCT116 cells (Figure 7). The first event induced by sister chromatid fusion is an anaphase bridge formation during the first mitosis (Figure 7-i). Such bridge would be resolved between anaphase and cytokinesis (Figur 7-ii), or persist in the next G1 phase (Figure 7-iii). The persisted chromatid bridge was proposed to cause cytokinesis furrow regression followed by daughter cell fusion (Figure 7-iv) [9] and/or be resolved by cytosolic nuclease because of abnormal nuclear envelope formation around the bridge (Figure 7-v) [10]. However, cell fusion was not associated with mCitrine expression in XpSIS cells (Figure 5A). In consistent with this observation, tetraploid cells did not increase upon sister chromatid fusion (Figure 3E and Supplementary figure 4A). The bridge resolution was shown to be associated with nuclear envelope rupturing [10], which is observable in our live cell imaging analysis because mCitrine harbors nuclear localization signal. However, we failed to observe the rupturing phenotype, possibly because of insufficient number of frames in the movies. Mitotic delay, which was observed in p53-compromised fibroblast cells upon TRF2 knockout [4], was also rare (Figure 7-vi)(Figure 5A). These results suggest that a single sister chromatid fusion was not enough to cause cell fusion and mitotic delay in HCT116 cells. We therefore assume that chromatin bridge in the next G1 was also eventually resolved (Figure 7-vii) presumably by some enzymatic activities [10]. The FISH analysis after reculturing of mCitrine positive XpSIS36-sgFUSION11 cells suggests that most surviving cells repaired broken bridge by either translocation or telomere healing by telomerase (Figure 7-viii, ix). A small fraction of cells possessed chromosome fusion with other chromosome, suggesting ongoing BFB cycle in these cells (Figure 7-x, xi). The increased aneuploidy of X chromosome at early time point indicates that fused sister X chromatids were segregated unequally in some cells (Figure 7-xii). Reduction of the cells carrying X aneuploidy in later time point suggests that aneuploid cells had reduced fitness and were removed from the population during long-term culturing (Figure 7-xiii) [38]. Finally, our live cell and statistical modeling indicate that MN was induced by a single X chromosome sister chromatid fusion in the first few cell cycle (Figure 7-xiv), and such MN formation possesses negative effect on the cellular fitness (Figure 7-xv). The formation of MN is consistent with a previous observation that about 23% of cells with H2B-GFP-visualized chromosome bridges during anaphase generated MN in the following G1 phase [39]. Such MN may be caused by abnormal elongation of the microtubule bundles bound to bridged chromosomes [40], or by enzymatic cutting of the bridge. Importantly, our FuVis-XpSIS system revealed that only one sister chromatid fusion could generate MN following the first few mitosis. The X chromosome fragmentation phenotype is consistent with reports that chromosomes in MN are prone to DNA damage during S phase due to abnormal nuclear envelope and subjected to fragmentation, which consequently gives rise to chromothripsis (Figure 7-xvi, xvii) [22] [41]. Such replication stress and DNA damage may contribute to cell cycle destabilization in MN positive cells. We therefore propose that, although most broken sister bridges can be repaired by either translocation or telomere healing and have minor effects on cellular fitness, even a single sister chromatid fusion can cause X chromosome missegregation, BFB cycle and/or MN formation followed by cell cycle destabilization, which potentially lead to chromosome rearrangements and tumorigenesis.

**Figure 7.**
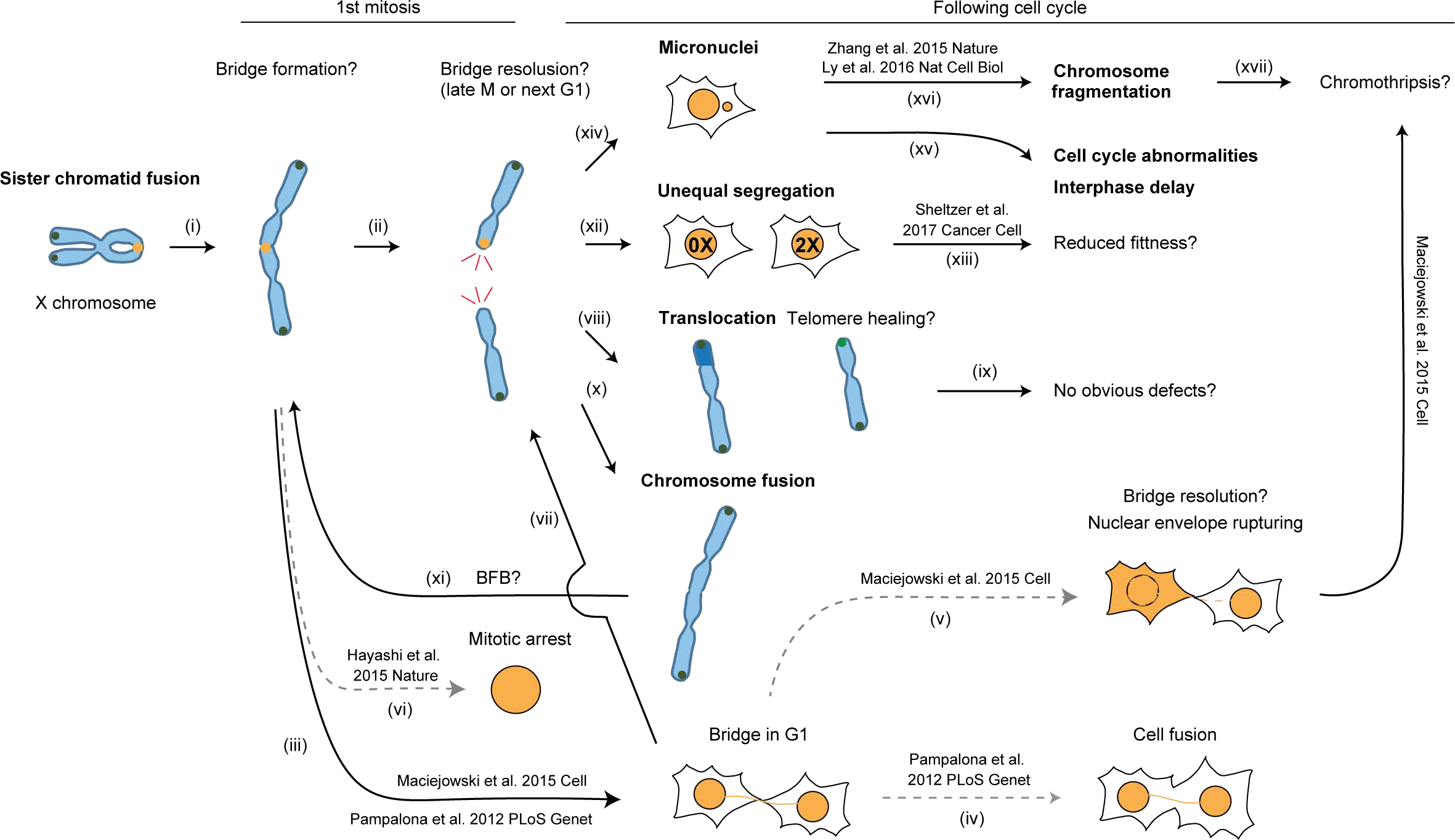
A model for the cellular fate of a single sister chromatid fusion. Read the main text for the detailed explanation. Bold characters indicate that these abnormalities were observed in FuVis-XpSIS system. Question mark indicates that the abnormalities were not directly observed but assumed from observed phenotypes. Dashed lines indicate that the abnormalities were observable but not observed in FuVis-XpSIS system.

### Perspective of FuVis

Fluorescent proteins have been widely applied to diverse studies from protein labelling to biological sensors [42]. We have expanded the applications of fluorescent proteins, namely a sensor of the specific chromosome rearrangement. The system described here can be applied to other types of rearrangement including specific translocations and other types of chromosome end-to-end fusion, such as inter-chromosome and intra-chromosome fusions (*i.e.* ring chromosome), all of which may contribute to chromosome-driven cellular transformation, tumor development and/or developmental disorders potentially through distinct mechanisms [1] [43]. Such expanded FuVis systems will provide unprecedented tools for the visualization and short- and long-term trace of specific chromosome rearrangements in user-defined cellular contexts, including mouse models and recently developing organoid models.

## Materials and Methods

### Cell culture

Human colon carcinoma HCT116 cells (ATCC) and their derivatives were cultured in DMEM (Nissui pharmaceutical) supplemented with 0.165% NaHCO3, 2 mM L-glutamine, 1 mM penicillin/streptomycine, 2.5 *µ*g mL^−1^ plasmocin (InvivoGen) and 10% fetal bovine serum. All cells were grown at 5% CO_2_ and ambient O_2_ condition. Cells were treated with 100 *µ*M *±*-blebbistatin (ab120425; Abcam) for 48 hr to induce tetraploidy.

### Plasmids

All plasmids used in this study are listed in supplementary file 4. For cloning of the sister cassette and control donor plasmid used for genomic integration, fragments of DNA that contains split mCitrine and potential CRISPR/Cas9 target sites were artificially synthesized (Eurofins Scientific) and used for the subsequent clonings. The potential CRISPR/Cas9 target sites were chosen from published non-targeting control sgRNA sequences that do not target human genome [44]. The neomycin resistant gene, 2× cHS4 insulator [29] and Xp subtelomere genomic sequence for homology template were added during cloning process. The resulting pMTH397 and pMTH729 were used for the generation of XpSIS and XpCTRL clones, respectively. DNA Sequences of pMTH397 and pMTH729 are provided in supplementary file 5 and 6, respectively. Sequence information of other plasmids are available upon request.

### CRISPR/Cas9-mediated homology-directed DNA cassette integration into genomic DNA

For DNA cassette integration, We used HCT116 cells because they possess relatively stable near-diploid chromosomes (n=45), highly efficient HR and carry only one X chromosome after losing Y chromosome [45] [46]. The CRISPR Design Tool (http://tools.genome-enineering.org) and the Cas-OFFinder (www.rgenome.net/cas-offinder/) were used to choose the target site for integration with minimum off-target sites. CRISPR/Cas9-based genome targeting was performed as described previously [47]. Briefly, HCT116 cells were plated in 6-well plate one day before transfection. The donor plasmids, pMTH397 or pMTH729, were transfected with eSpCas9(1.1)-sgCHRXpYp-Subtel2 (pMTH393) that targets chromosome Xp subtelomere locus using FuGENE HD reagent (Promega). Two days post transfection, cells were diluted and plated on 10 cm dish with 700 *µ*g mL^−1^ G418. Medium with G418 was refreshed every 3 days for 2 weeks, and individual colonies were isolated. Genomic DNA were obtained and genomic integration was assessed by PCR using primers listed in supplementary file 7 and southern blotting as described below. The PCR products were sequenced to confirm integration at the expected locus.

### Southern blotting

Genomic DNA was digested with EcoRI, run on 0.7% SeaKem GTG agarose (Lonza), and transferred to the Amersham Hybond-N+ membrane (GE Healthcare Life Sciences). For the mCit-C probe, an 833 bp fragment was amplified by PCR using MTH384 and MTH417 as primers and pMTH393 as a template. The probe was generated by random labeling and hybridized to the membrane at 63°C.

### Viral infection

The lentivirus particles were generated as described previously [48] with minor modifications. Briefly, HEK293FT cells (Thermo Fisher Scientific) were transfected with transfer plasmid, psPAX2 (a gift from Didier Trono, addgene #12259) and pCMV-VSV-G (a gift from Bob Weinberg, addgene #8454) using polyethyenimine (PEI). Medium was replaced on the next day, and medium containing active lentivirus particles was collected on day 2 and day 3 post transfection. For LentiCRISPR-sgEMPTY and LentiCRISPR-sgFUSIONs, cells were infected in growth media containing 8 *µ*g mL^−1^ polybrene and lentivirus, and cultured for 2 days. Puromycin was added to the culture at 1 *µ*g mL^−1^ and infected cells were selected for more than 2 days before analysis. The amount of lentivirus required for nearly 100% infection was determined empirically. All target sequences of CRISPR/Cas9 used in this study are listed in supplementary file 1.

### Fluorescent in situ hybridization

Conventional fluorescent in situ hybridization was performed as described previously [49] with modifications as described below. Briefly, cells were exposed to 100 ng mL^−1^ colcemid for 2 hr and fixed in 3:1 Methanol/Acetic acid for 6 minutes. For telomere and centromere double-staining, FAM-conjugated telomeric PNA probe (TelC-FAM; Panagene) and Cy3-conjugated centromeric PNA probe (Cent-Cy3; Panagene) were used. For telomere, chromosome X centromere, and sister DNA cassette triple-staining, TelC-FAM, Dig-conjugated chromosome X centromere specific probe (HXO-10 X:Dig; Chromosome Science Lab), and Cy5-labeled pMTH368 probe (on-demand probe; Chromosome Science Lab) were mixed in PNA hybridization solution (70% (v/v) formamide, 0.25% (w/v) Blocking Reagent, 10 mM Tris-HCl pH7.5) and used for hybridization at 70°C for 5 min. After overnight incubation at 37°C, slides were washed in 2× SSC for 5 min at RT, 50% formamide/2× SSC for 20 min at 37°C, and 1× SSC for 15 min. The same hybridization and washing protocol was used for chromosome X centromere and control DNA cassette double-staining with green fluorophore-labeled chromosome X centromere probe (XHO-10 X:Green; Chromosome Science Lab) and Cy3-labled pMTH727 probe (on-demand probe; Chromosome Science Lab). For X chromosome and chromosome X centromere double-staining, orange fluorophore-conjugated X chromosome painting probe (XCP X orange; Metasystems) and green fluorophore-conjugated chromosome X centromere and orange fluorophore-conjugated chromosome Y centromere specific probe (XCE X/Y; Metasystems) were used according to the manufacture’s instruction. Since male-derived HCT116 cells lost Y chromosome, XCE X/Y probe did not give any orange signal. In the structural abnormality analysis, inter-chromosome fusion was distinguished from translocation by the presence of narrow chromosome region in non-X chromosome, which suggest the presence of centromere.

### Flow cytometry

Cells were harvested by trypsinization, resuspended in 1×PBS with 0.1 mM EDTA and filtered through 5 ml polystyrene round-bottom tube with cell-strainer cap (Corning) before running through the FACSAria III flow cytometer/cell sorter (Becton Dickinson). Dead cells were excluded by positive PI-staining, and single cells were gated by low FSC-W value before analysis and sorting. mCitrine positive cells were detected by 488 nm laser and 530/30 filter set.

### Fusion junction analysis

For sister chromatid fusion junction analysis, mCitrine positive XpSIS15-sgFUSION11, XpSIS15-sgFUSION4, and XpSIS36-sgFUSION11 cells were sorted by FACSAria III at 6 days post infection and incubated for another 11 days before extraction of genomic DNA. The genomic DNA was used as a template for PCR using MTH672 and MTH673 as primers. PCR products were gel-purified and cloned into EcoRV site of pBSII plasmid by in-fusion reaction, followed by sequencing of individual clones. For sister cassette repair junction analysis, XpSIS15-sgFUSION11 cells were harvested at 15 days post infection and the genomic DNA was used for PCR with primer set MTH672 and MTH803. For control cassette repair junction analysis, XpCTRL48-sgFUSION11 cells were harvested at 10 days post infection and the genomic DNA was used for PCR with primer set MTH672 and MTH806. All primer sequences are listed in supplementary file 7.

### Live-cell imaging

Live-cell imaging was performed in conventional cell culture dishes or plates placed on a BZ-X710 all-in-one fluorescence microscope (KEYENCE) equipped with stage-top chamber and temperature controller with built-in CO_2_ gas mixer (INUG2-KIW; Tokai hit) and a 10X 0.3 NA air CFI Plan Fluor DL objective (Nikon) at 37 °C and 5% CO_2_. mCitrine expression was detected by metal-halide lamp and YFP-optimized filter cube (ex: 500/10 nm, em: 535/15 nm, dichroic: 515LP)(M square). Images were captured by a BZ-H3XT time-lapse module typically every 6 to 12 minutes for at least 66 hours. The fate of mCitrine positive cells were observed manually by eyes. For all mCitrine positive cells, the beginning time of mCitrine expression, the time of nuclear envelope break down and cytokinesis, and the time of abnormalities including cell fusion, cell death, mitotic slippage and tripolar mitosis were recorded. MN or multi-nuclei including binuclei phenotype were determined by mCitrine localization. For XpSIS cells, the time when mCitrine becomes too dim to observe was also recorded. The fates of mCitrine positive cells for all different lineages were recorded as a tidy data (supplementary file 8, 9) and used for the statistical analysis and lineage tree visualization.

### Statistical analysis

We adopted ‘high-level descriptive (top-down) statistical models’ [50] to the data of live cell imaging. In the analysis, each lineage has a group of cells and it would be natural to assume that the cells share some common background characteristics affecting the observations such as the MN formation and interphase duration. If we ignore the cluster structure and consider each observation for each cell as a random variable subjected to an independent and identical distribution, that would lead to a bias to the interpretation of the data [51]. Therefore, we explicitly implement the clustered or hierarchical structure into the statistical models. Furthermore, we constructed the alternative non-hierarchical model as well as other possible alternatives. We applied the models to the data and built predictive distributions on each model to make them approximate to the unknown distribution that generated the data such as whether MN was formed or not (*MN_n_*, in the Figure 5C). The models and predictive distributions were defined and implemented by a probabilistic programming language Stan [52]. We computed through the package ‘rstan’ in the statistical computing environment R [53]. In the programs, the Widely Applicable Information Criterion (WAIC) values were calculated to evaluate the appropriateness of the predictive distributions [23] [24]. It should be noted that the assessment of the model is from the point of view of ‘prediction’ originally proposed in the theory of AIC (Akaike Information Criteriorn) [54] [55] and that WAIC is an extended version of AIC in a Bayesian framework. WAIC is applicable to the models that implement parameters whose posterior distribution does not resemble any normal distribution. The details of the models and the process of the data analysis is written in the supplementary file 3. For cell cycle abnormality analysis in Figure 6A, we built GLMs, performed the maximum likelihood estimation and calculated AIC values (Supplementary file 3, 10, 11).

### Lineage tree visualization

The tidy data was used to visualize lineage trees of all cell lineages analyzed in the live cell imaging (supplementary file 8, 9). The code is available in the supplementary file 12. The trees of the following lineages were manually modified with Adobe Illustrator CC 2019: 22, 23, 53, 54, 60, 61, 137, 138, 167, 185, 186, 197, 467, 550, 562, 597, and 598.

## Supporting Information

All supplementary files are found at https://figshare.com/s/3fd3fb4e5571f14f1ecf.

## Acknowledgments

We thank Robert Weinberg, Didier Trono, Feng Zhang and Itaru Imayoshi for sharing plasmids, Masato Kanemaki for sharing methods, Diana Zamora Romero and Hiroya Fukuda for experimental supports with cloning and/or sequencing, Mari Tsuboi for her assistance with cloning and data acquisition from live cell imaging, Hisao Masukata for critical reading of the manuscript and Ishikawa lab members for helpful discussion. Makoto T Hayashi is supported by grants from the Senri Life Science Foundation, the Uehara Memorial Foundation, the Daiichi Sankyo Foundation of Life Science, the Nakajima Foundation, The Mochida Memorial Foundation Research Grant, Grant-in-Aid for Young Scientists (A) (16H06176), Grant-in-Aid for Scientific Research on Innovative Areas (16H01406), and the Kyoto University Hakubi project.

