## Supplementary Figures for "A Single Defined Sister Chromatid Fusion Destabilizes Cell Cycle through Micronuclei Formation"

**Supplementary Figure 1. Validation of XpSIS clones. A**, Schematic representing a possible outcome of CRISPR/Cas9-targeting of subtelomere regions. Digestion of two different subtelomeres in the G1 phase can result in inter-chromosomal fusion. Digestion of two different subtelomeres in the G2 phase results in either inter-chromosome-type fusion or inter-chromatid type fusion, while targeting of sister chromatid subtelomeres can generate sister chromatid fusion. **B**, FISH images of sister chromatid fusion and chromosome-type fusion induced by CRISPR/Cas9-metiated digestion of multiple subtelomeres in IMR-90 E6E7 cells. Representative images show centromere (red), telomere (green) and DAPI (blue). Triangles in enlarged images indicate sister chromatid fusion (top) and inter-chromosome fusion (bottom). Scale bar, 10 $\mu$m. **C**, Schematic of PCR to screen the sister cassette integration at Xp subtelomere. MTH632, MTH653 and MTH698 are primers used for genomic PCR. The shorter fragment (1,188 bp) was used as a positive control. **D**, Representative results of PCR for screening FuVis-XpSIS clones1 and 2. **E**, Bar graph shows percentage of mCitrine positive cells. Indicated clones were infected with lentivirus carrying CRISPR/Cas9-sgFUSION4 and analyzed at 6 days post infection. **F**, Bar graph shows percentage of mCitrine positive cells. Indicated clones were infected with lentivirus carrying CRISPR/Cas9-sgFUSION11 and analyzed at 10 days post infection. **G**, Schematics of Southern hybridization to confirm the sister cassette integration at the single locus. **H**, Southern hybridization result using mCit-C probe in XpSIS15 and XpSIS36 cells. EtBr indicates total genomic DNA digested with EcoRI. **I**, Percentage of mCitrine positive XpSIS15 and XpSIS36 cells upon CRISPR/Cas9-targeting of multiple different loci across the spacer and the neoR gene. The map under the graph represents location of the target sites.

**Supplementary Figure 2. Sequence analysis of fusion junction in XpSIS. A**, **B**, Representative FISH images of FuVis-XpSIS15-sgEMPTY (A) and -sgFUSION11 (B) cells showing the sister cassette (red), telomere (green) and DAPI (blue). Whole cells and mCitrine positive cells were harvested (A) and sorted (B), respectively, at 8 days post infection. Bar, 10 $\mu$m. **C**, Schematic representing PCR product used for fusion junction analysis. **D**, Result of PCR with the fusion primer set in (C) using indicated genomic DNA. The product specific in sgFUSION11 condition was cloned into pBSII vector and sequenced. **E**, Schematic representing PCR product used for repair junction analysis. **F**, Percentage of truncated features at fusion junctions in XpSIS36-sgFUSION11 and XpSIS15-sgFUSION4. **G**, **H**, Distribution of truncated length at fusion junctions in XpSIS36-sgFUSION11 (G) and XpSIS15-sgFUSION4 (H). **I**, Percentage of junction features in XpSIS36-sgFUSION11 and XpSIS15-sgFUSION4. Complex indicates junctions with multiple independent integration of the original cassette sequences.

**Supplementary Figure 3. Validation of XpCTRL48 and characterization of FuVis-Xps.** **A**, Schematic of PCR to screen the control cassette integration at Xp subtelomere. MTH442, MTH653, and MTH698 are primers used for genomic PCR. The shorter fragment (1,188 bp) was used as a positive control. **B**, Representative result of PCR for screening FuVis-XpCTRL clones. Clone48 has duplication of homology arm. **C**, Schematics of Southern hybridization to confirm the control cassette integration at the single locus. **D**, Southern hybridization result using mCit-C probe in XpCTRL48 cells. EtBr indicates total genomic DNA digested with EcoRI. **E**, Representative FISH images of FuVis-XpCTRL48 cells showing the control cassette (red), chrX centromere (green) and DAPI (blue). Bar, 10 $\mu$m. **F**, **G**, Schematics showing the fate of mCitrine gene after sister chromatid fusion in FuVis-XpSIS (F) and repair in FuVis-XpCTRL (G). **H**, Schematic of flow cytometery analysis. FuVis-Xps cells infected with CRISPR/Cas9-sgFUSION11 lentivirus were selected with puromycin. mCitrine-positive cells were sorted at 8 days post infection, re-cultured for 13 days and analyzed by flow cytometery. **I**, Flow cytometery analysis as described in (H). Square indicates mCitrine-positive population in each condition. **J**, Bar graph representing average percentage of mCitrine-positive cells after re-culturing. Dot plots indicate individual data (n = 3).

**Supplementary Figure 4. Analysis of chromosome spread in XpCTRL48-sgFUSION11 and XpSIS36-sgFUSION11.** **A**, The number of total chromosomes in indicated cells. Counts of 90 spreads from three independent experiments are shown (n=30 per experiment). Dotted lines indicate 45, 90 and 135, which represent near-diploid, near-tetraploid, and near-octaploid cell, respectively. **B**, FISH images of FuVis-XpSIS36-sgFUSION11 D8 cells as in Figure 3B-D.

**Supplementary Figure 5. Lineage trees of FuVis-XpCTRL48-mCit (N+n)**. Symbols representing cell cycle progression and abnormalities are summarized in Figure 4C.

**Supplementary Figure 6. Lineage trees of FuVis-XpCTRL48-sgFUSION11 (1+n).** Symbols representing cell cycle progression and abnormalities are summarized in Figure 4C.

**Supplementary Figure 7. Lineage trees of FuVis-XpCTRL48-sgFUSION11 (N+n).** Symbols representing cell cycle progression and abnormalities are summarized in Figure 4C.

**Supplementary Figure 8. Lineage trees of FuVis-XpSIS15-mCit (N+n).** Symbols representing cell cycle progression and abnormalities are summarized in Figure 4C.

**Supplementary Figure 9. Lineage trees of FuVis-XpSIS15-sgFUSION11 (1+n).** Symbols representing cell cycle progression and abnormalities are summarized in Figure 4C.

**Supplementary Figure 10. Lineage trees of FuVis-XpSIS15-sgFUSION11 (N+n).** Symbols representing cell cycle progression and abnormalities are summarized in Figure 4C.

**Supplementary Figure 11. Lineage trees of FuVis-XpSIS36-mCit (N+n).** Symbols representing cell cycle progression and abnormalities are summarized in Figure 4C.

**Supplementary Figure 12. Lineage trees of FuVis-XpSIS36-sgFUSION11 (1+n).** Symbols representing cell cycle progression and abnormalities are summarized in Figure 4C.

**Supplementary Figure 13. Lineage trees of FuVis-XpSIS36-sgFUSION11 (N+n).** Symbols representing cell cycle progression and abnormalities are summarized in Figure 4C.

**Supplementary Figure 14. Lineage analysis of mCitrine positive cells. A**, Table representing the numbers of lineages that possess indicated abnormalities in indicated conditions. The numbers were used for calculation of the percentages of lineages shown in Figure 5A. The same lineage was counted more than twice if multiple abnormalities were observed in the lineage. **B**, Representative live cell images of FuVis-XpSIS36-sgFUSION11 cell that possessed MN formation following the first (top) and second (bottom) mitosis upon sister chromatid fusion. Yellow triangle, the beginning of mCitrine expression. White triangle, MN. ID, lineage ID. Bar, 10 $\mu$m. **C**, Lineages that show MN formation during 1+n cell cycle in XpSIS15-sgFUSION11. The numbers above the trees represent lineage ID. **D**, Posterior distribution of the parameters inferred from the second predictable model 1_3 with median (black circle), and 50% (red bar) and 95% (black bar) credible intervals. **E**, Dot plots representing average posterior predicted probabilities of MN formation calculated by using the parameters inferred in the model 1_3.

**Supplementary Figure 15. Modeling of the effect of MN on interphase duration. A**, Six model structures constructed to explain interphase duration ($Int\_duration_n$), which is subjected to exponential log distribution (top). The $Int\_duration_n$ is modeled by the following function and parameters: LogNormal, exponential log function; *micro*, the coefficient of MN; *scf*, the coefficient of sister chromatid fusion; *rnf*, the coefficient of repair (no fusion); *stg*, the coefficient of cell cycle stage after mCitrine expression (1+n or N+n); *sis15* and *sis36*, the coefficient of cell line (SIS15 or SIS36 compared to CTRL48); $lin_i$, *i=1,...,N_{lineages}*, intercepts representing individuality (unknown cellular characteristics shared in each lineage); and *b*, a bias parameter. Large capitals indicate variables (dummy variables) obtained from data (0 or 1 in supplementary file 8). The models implement all explanatory variables with individuality (2_1), all explanatory variables without individuality (2_2), explanatory variables without MN and individuality (2_3), only MN, SCF and individuality (2_4), only individuality (2_5), and only MN and SCF without individuality (2_6). The calculated WAIC values for each model are shown on right. **B**, Causal diagram that we assume for the inference of the impact of MN on interphase duration. Experimentally controlled variables, SCF, SIS, RNF, and Stage_N are given upstream variables (confounding factors), thereby they are added to the linear predictor in order to fix them and infer the causality between MN and interphase duration. **C**, Posterior distribution of the indicated parameters inferred from the most predictable model 2_2 with median (black circle), and 50% (red bar) and 95% (black bar) credible intervals.

Supplementary figure 1 Kagaya et al.

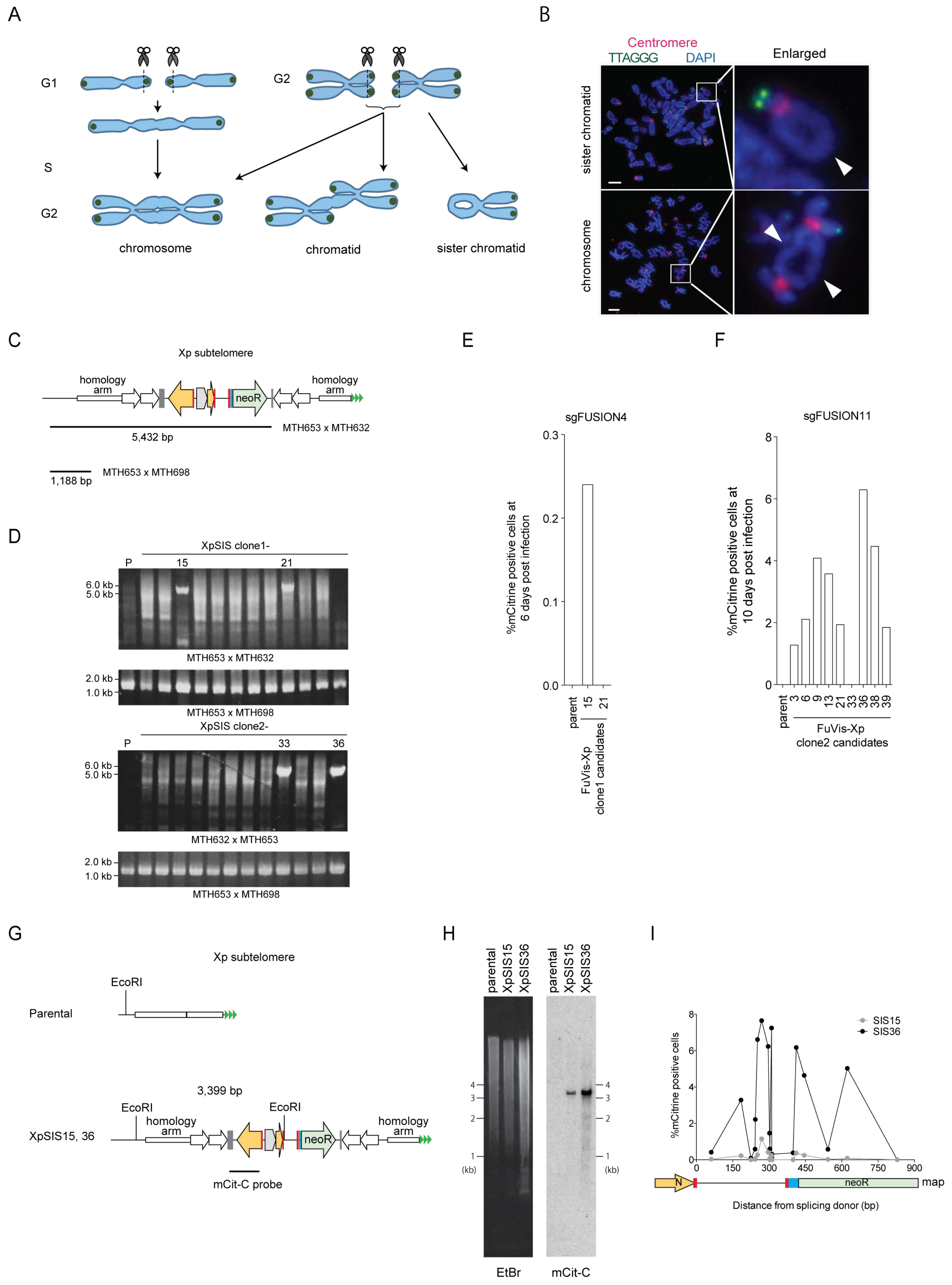

Supplementary figure 2 Kagaya et al.

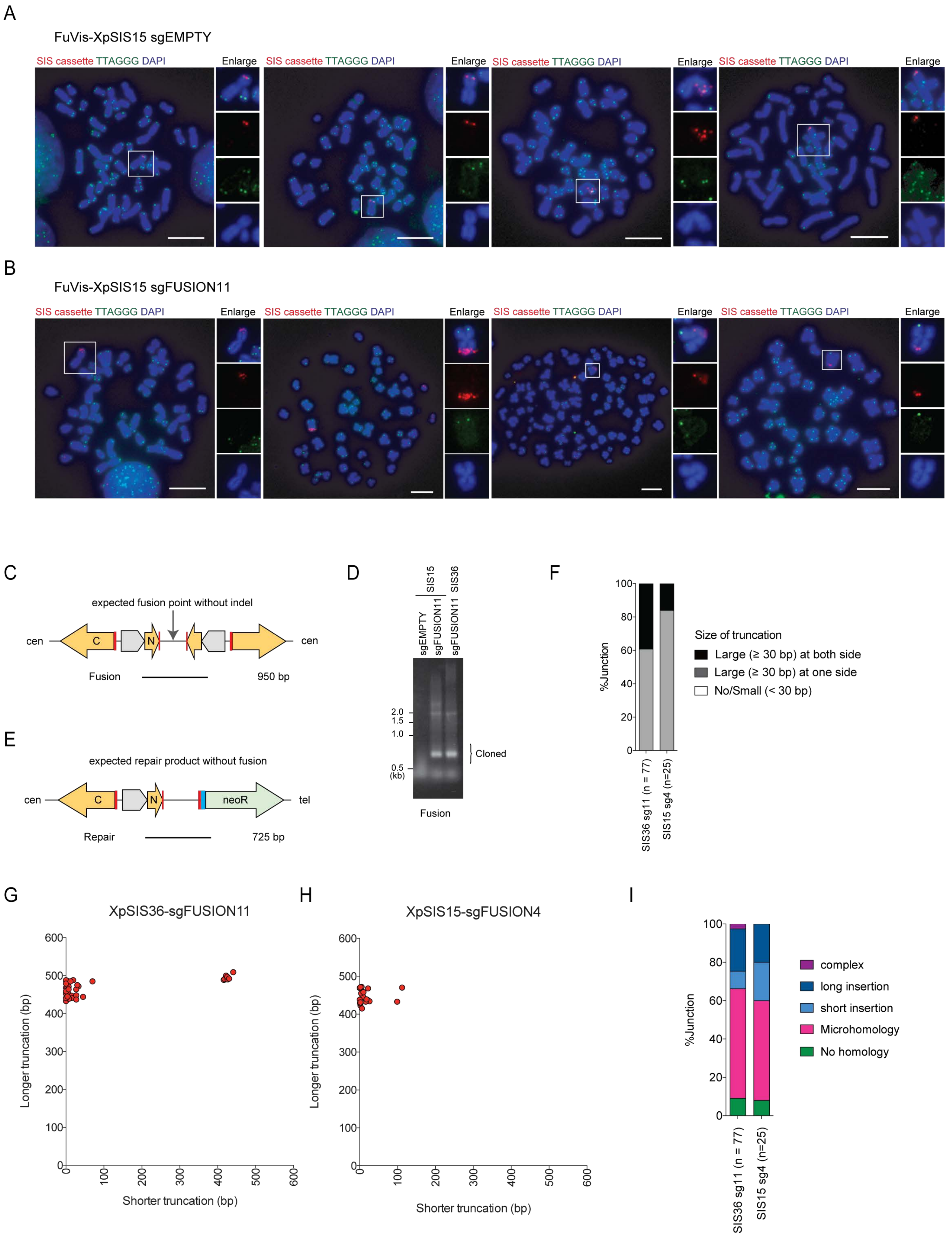

Supplementary figure 3  Kagaya et al.

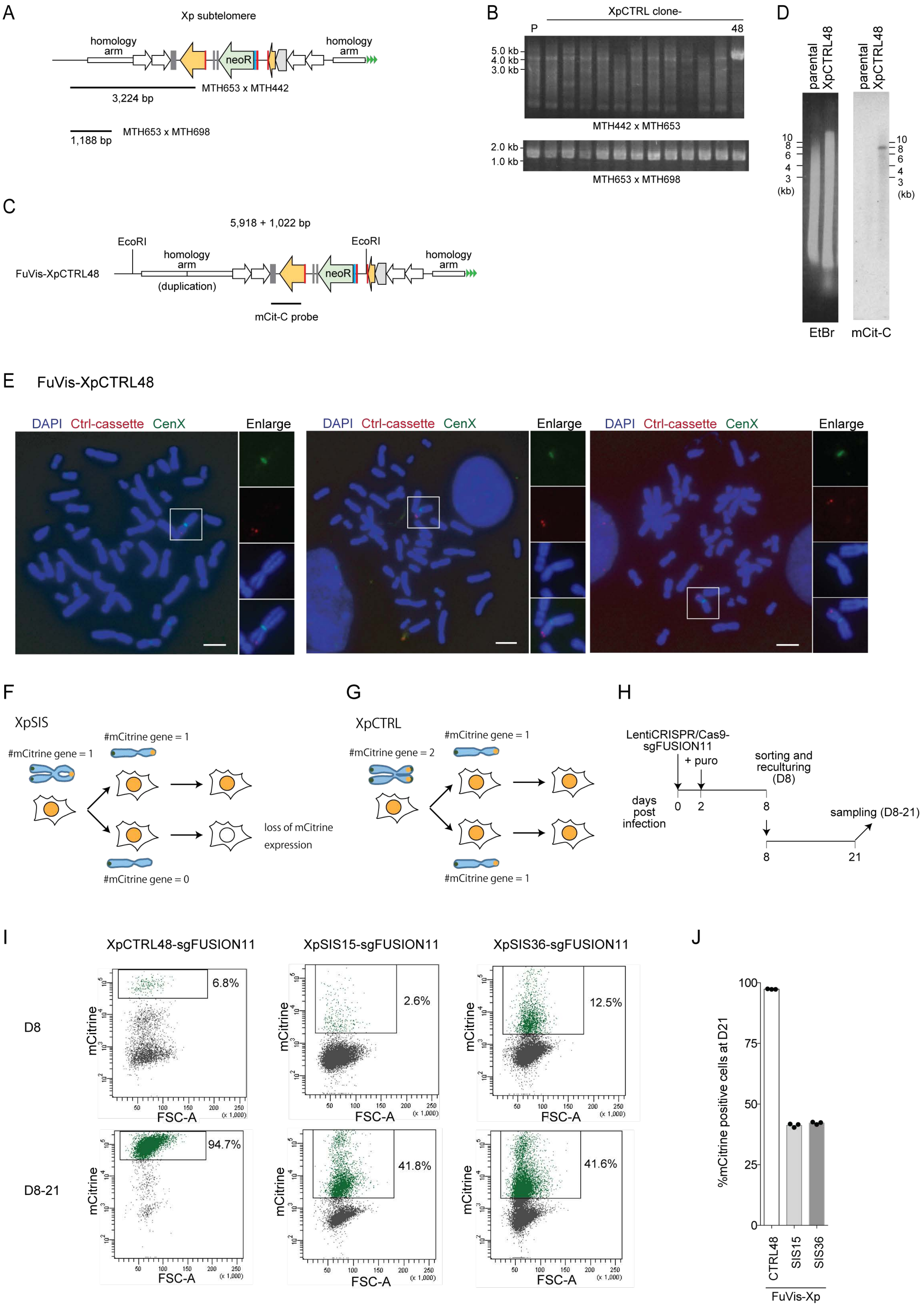

Supplementary figure 4  Kagaya et al.

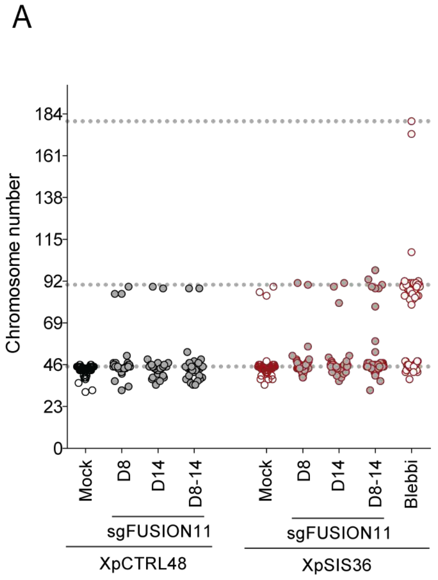


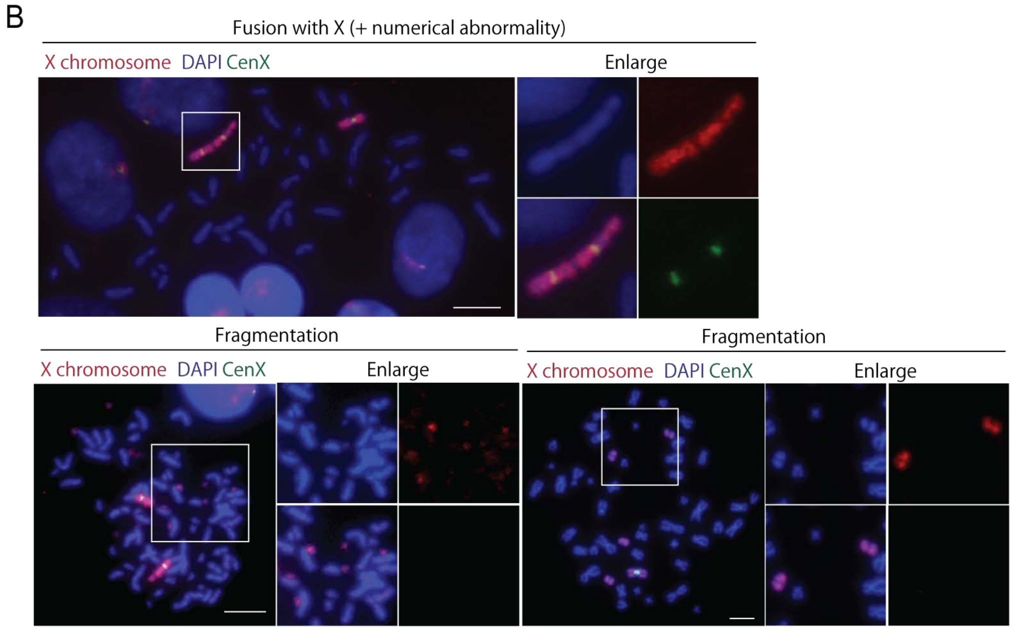

Supplementary figure 5 Kagaya et al.

### FuVis-XpCTRL48 pMX-mCitrine

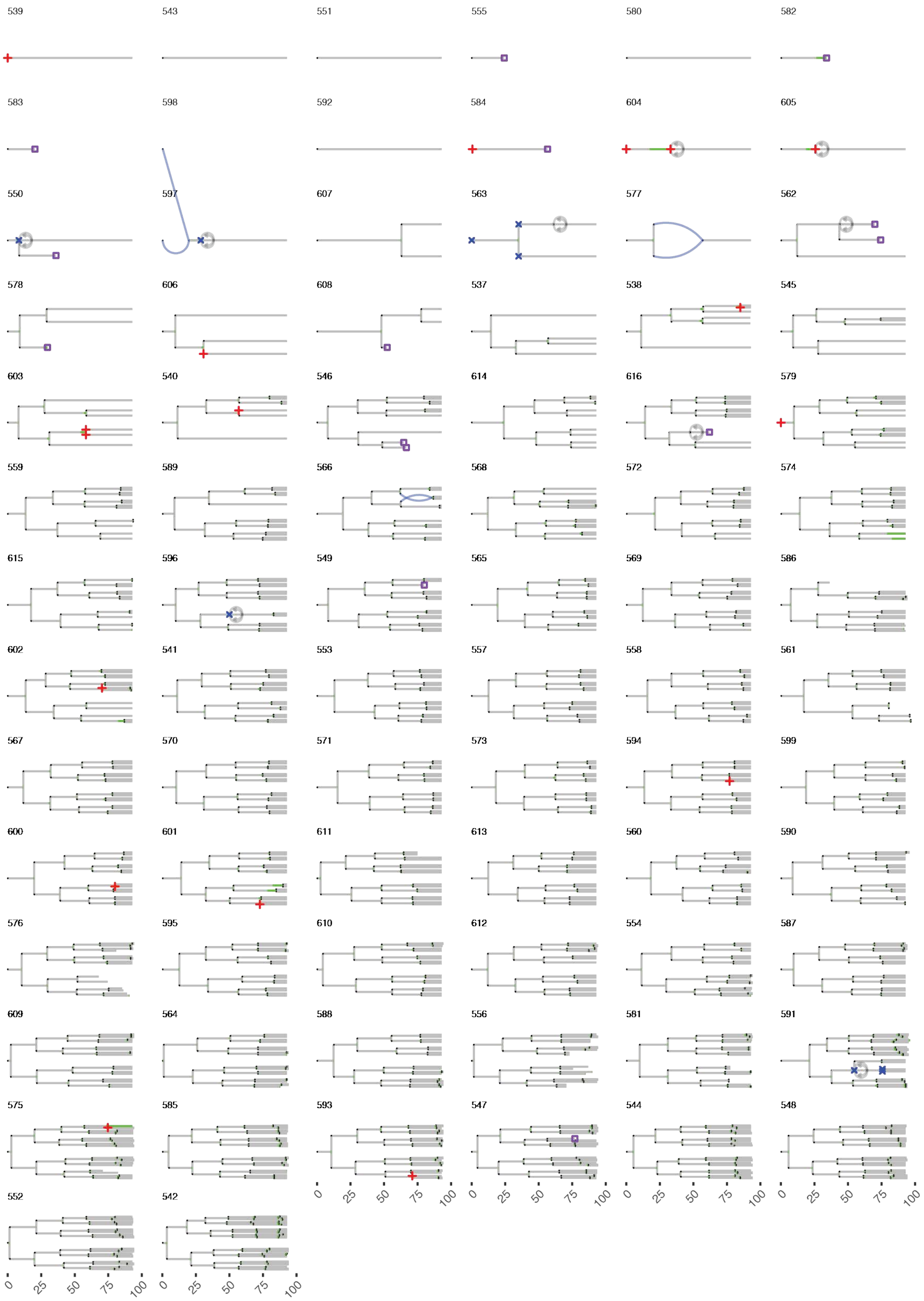

Supplementary figure 6 Kagaya et al.

FuVis-XpCTRL48 LentiCRISPR/Cas9-sgFUSION11 1+n cell cycle

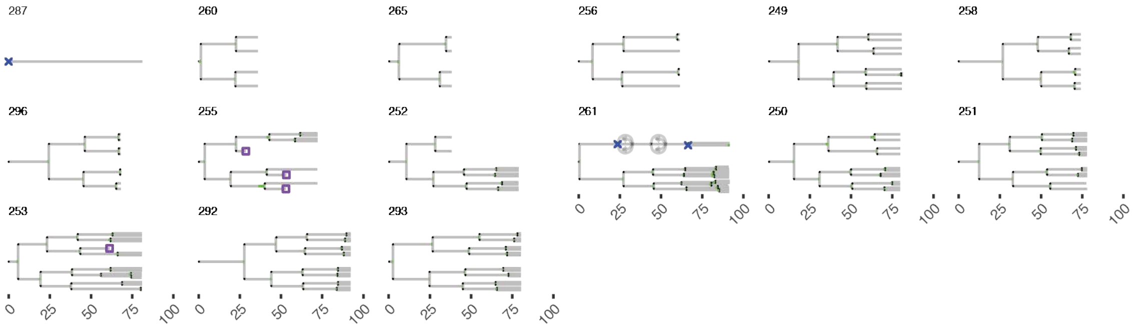

Supplementary figure 7 Kagaya et al.

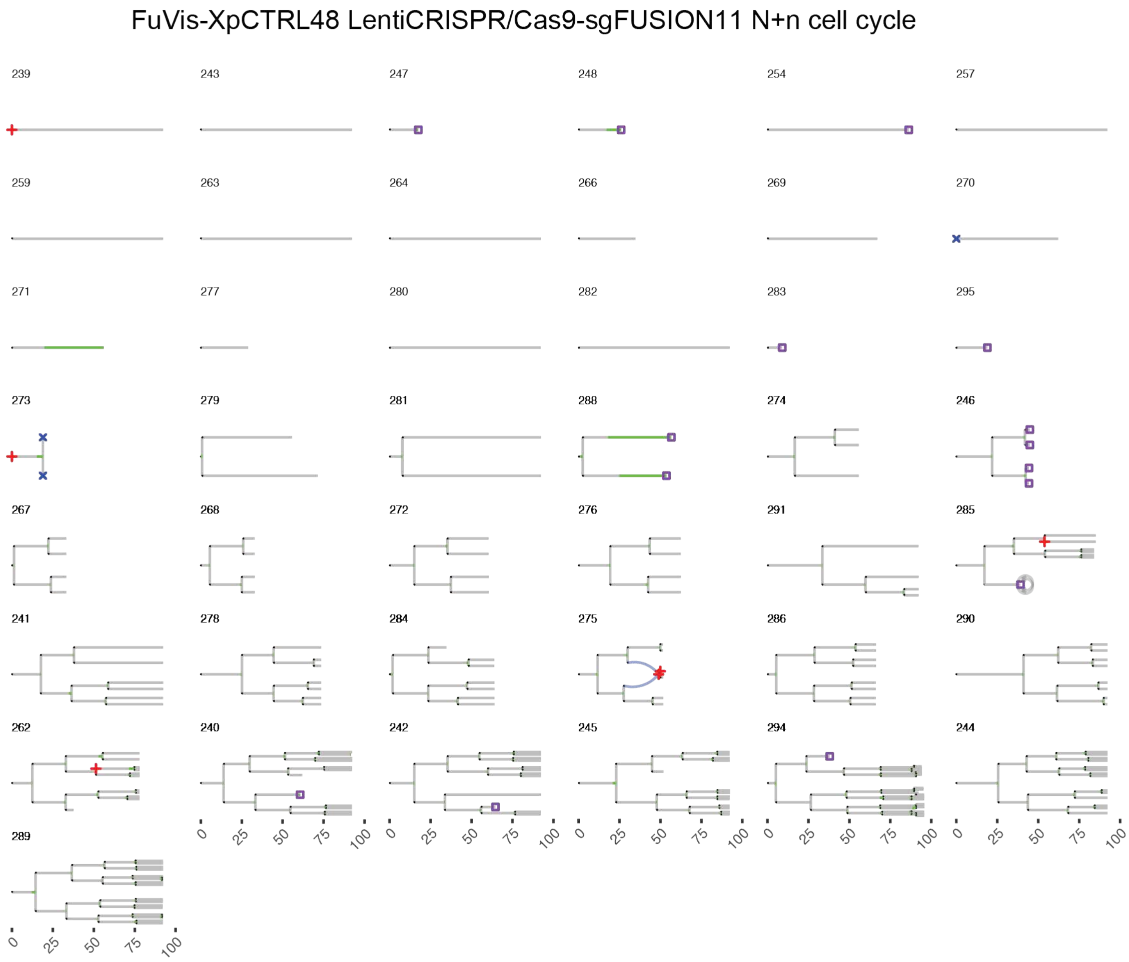

Supplementary figure 8 Kagaya et al.

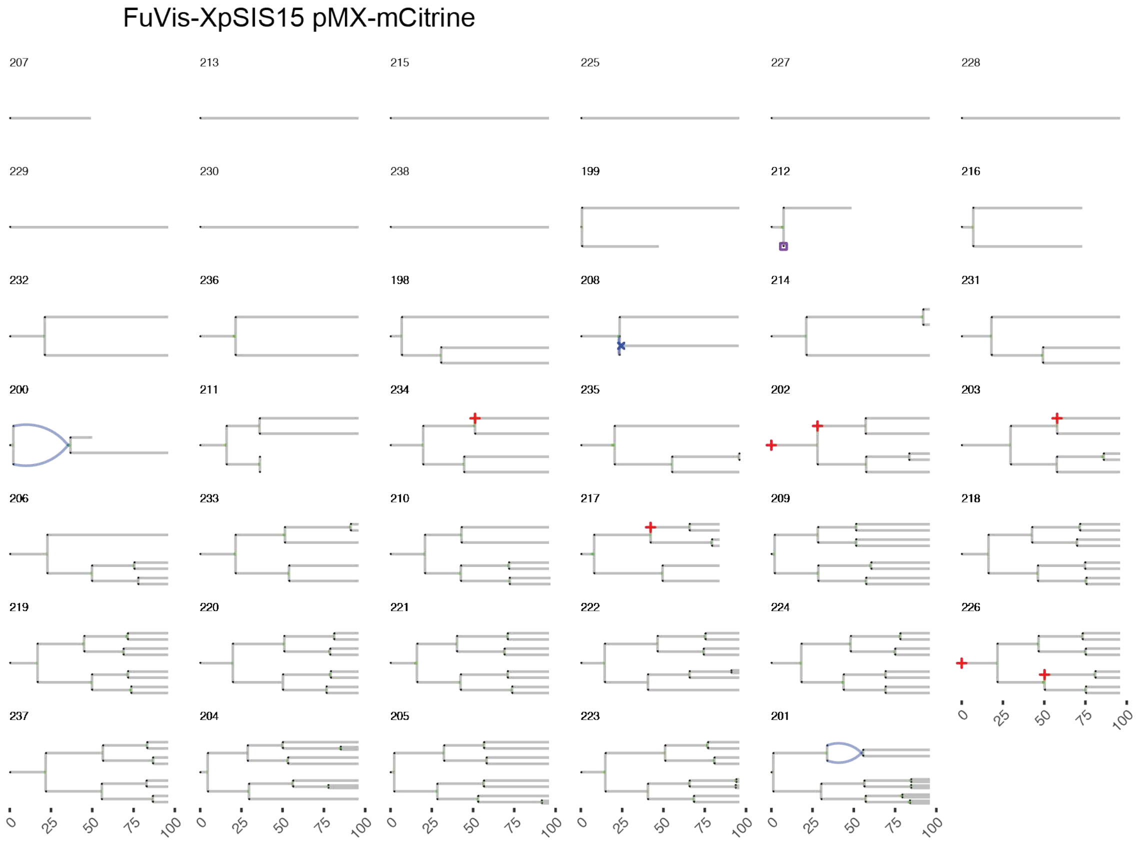

Supplementary figure 9 Kagaya et al.

FuVis-XpSIS15 LentiCRISPR/Cas9-sgFUSION11 1+n cell cycle

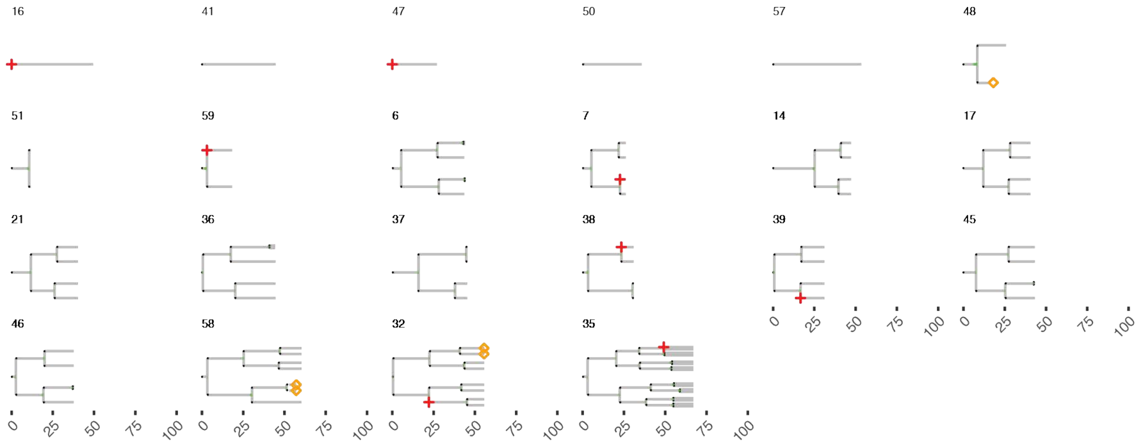

Supplementary figure 10 Kagaya et al.

FuVis-XpSIS15 LentiCRISPR/Cas9-sgFUSION11 N+n cell cycle

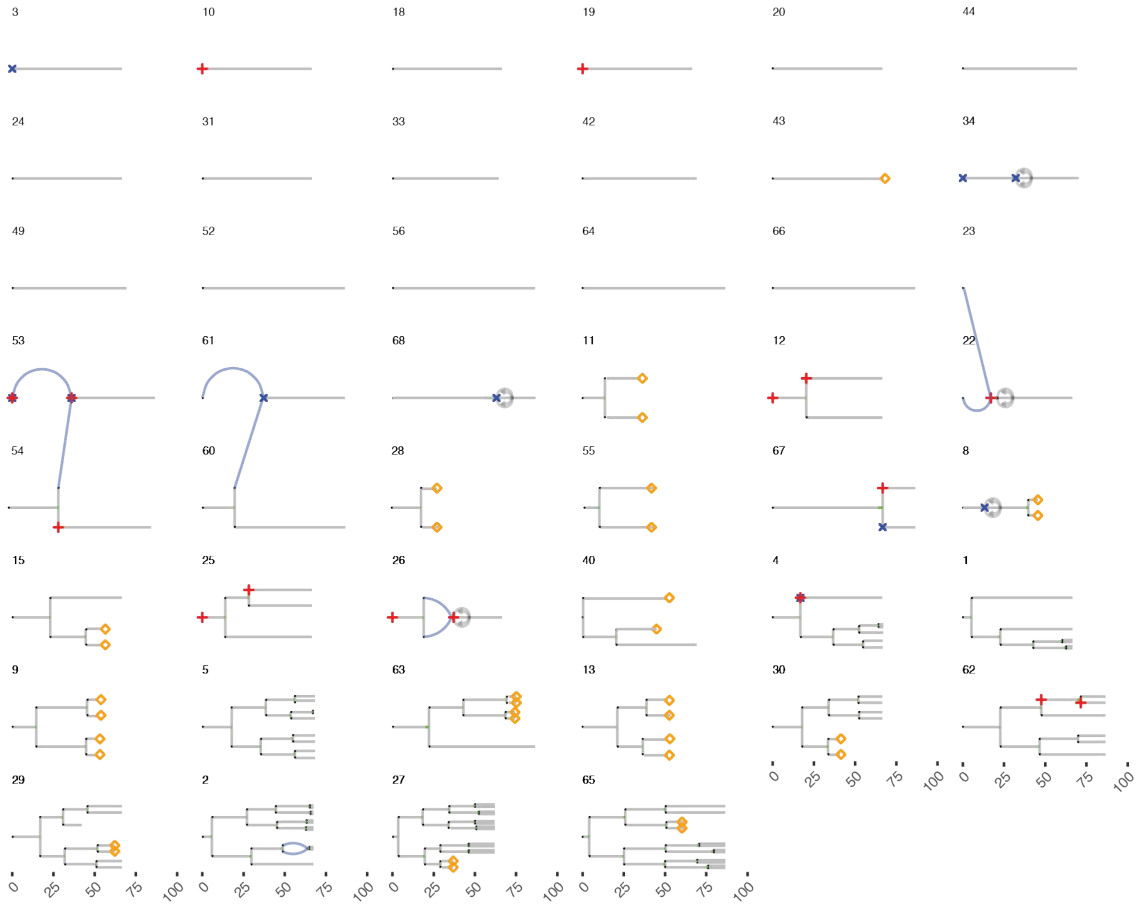

Supplementary figure 11 Kagaya et al.

FuVis-XpSIS36 pMX-mCitrine

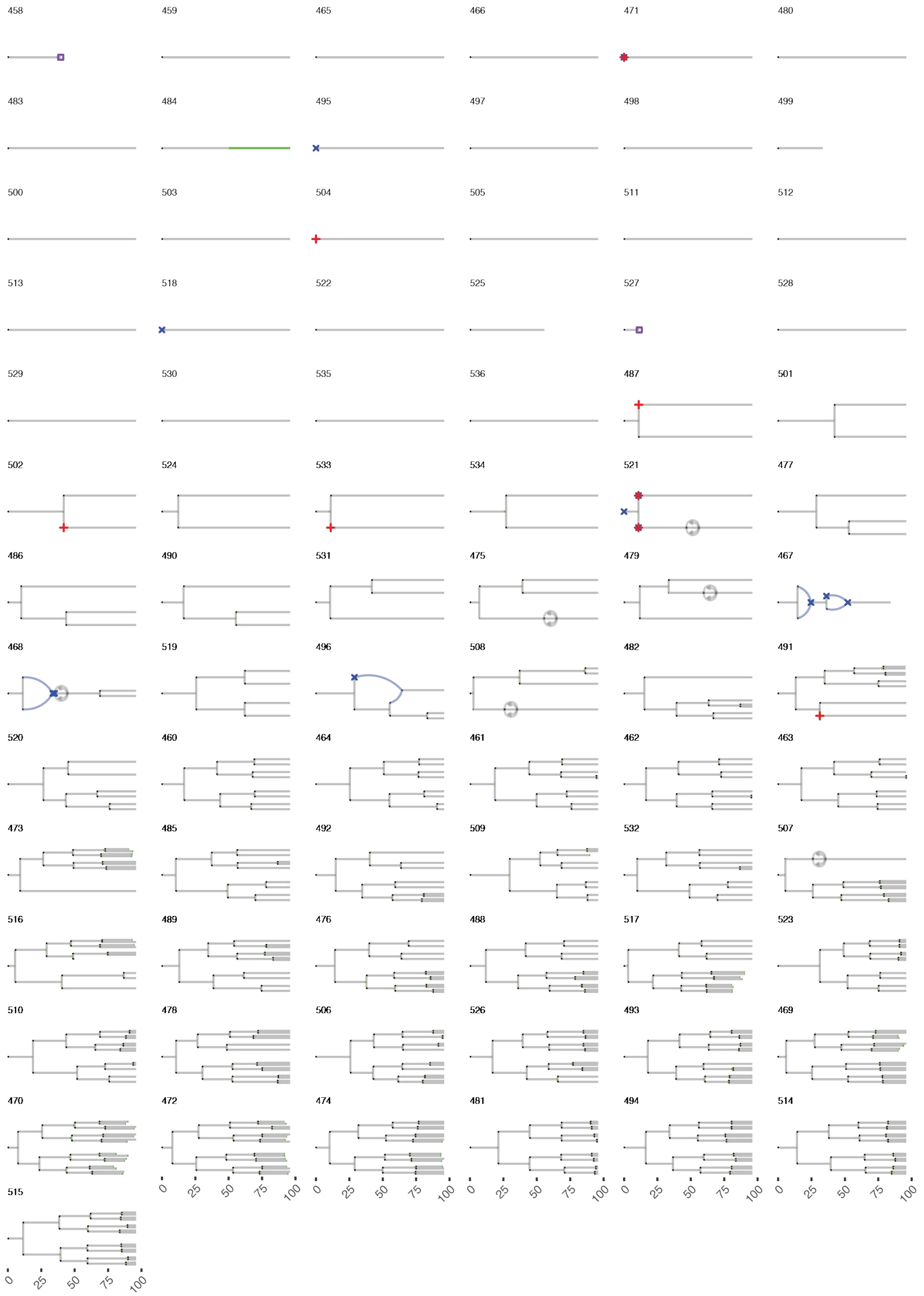

Supplementary figure 12 Kagaya et al.

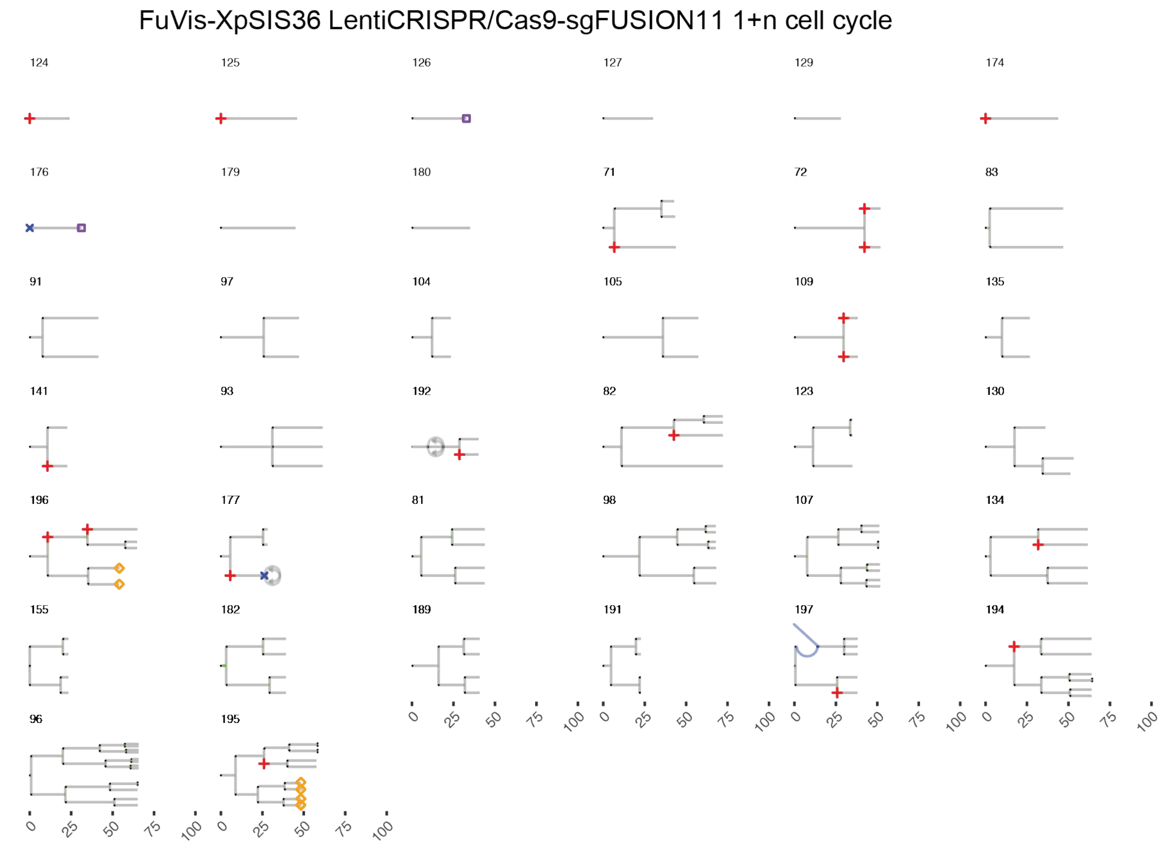

Supplementary figure 13 Kagaya et al.

FuVis-XpSIS36 LentiCRISPR/Cas9-sgFUSION11 N+n cell cycle

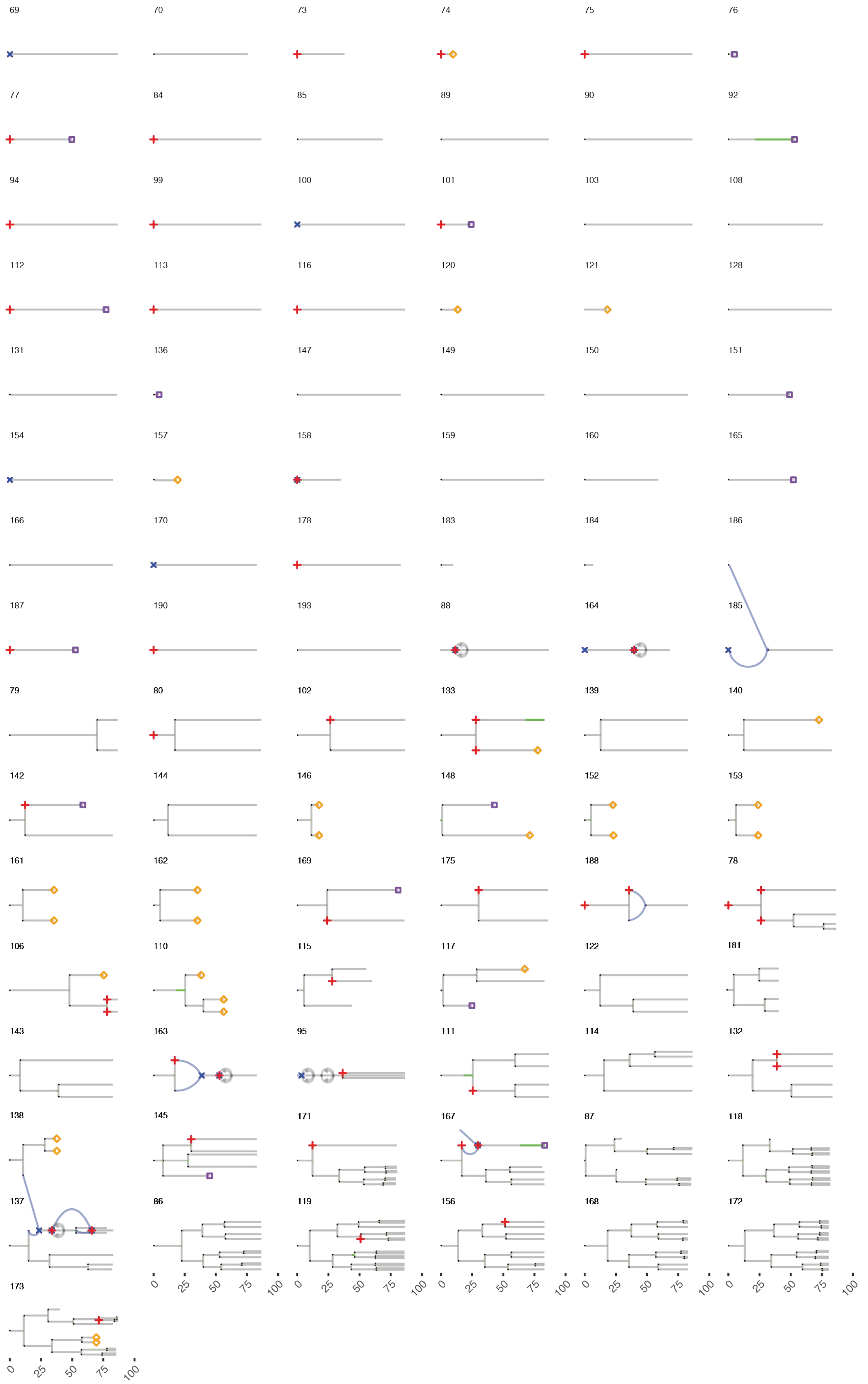

Supplementary figure 14  Kagaya et al.

A

| | | | Number of lineages that possess indicated abnormalities | | | | | | | | |
|---|---|---|---|---|---|---|---|---|---|---|---|
| | | | No mitosis | Mitotic delay | Micronuc | Bi/multinuc | Cell death | Regress | Cell fusion | Fading | Total |
| CTRL48 | mCit | N+n | 9 | 12 | 15 | 5 | 12 | 9 | 4 | 0 | 80 |
| | sgFUSION11 | 1+n | 1 | 2 | 0 | 2 | 2 | 1 | 0 | 0 | 15 |
| | | N+n | 12 | 6 | 5 | 2 | 11 | 1 | 1 | 0 | 43 |
| SIS15 | mCit | N+n | 9 | 0 | 5 | 1 | 1 | 0 | 2 | 0 | 41 |
| | sgFUSION11 | 1+n | 5 | 1 | 8 | 0 | 0 | 0 | 0 | 3 | 22 |
| | | N+n | 15 | 5 | 11 | 7 | 0 | 5 | 8 | 14 | 52 |
| SIS36 | mCit | N+n | 25 | 1 | 7 | 5 | 2 | 5 | 3 | 0 | 79 |
| | sgFUSION11 | 1+n | 7 | 1 | 15 | 2 | 2 | 2 | 0 | 2 | 38 |
| | | N+n | 29 | 5 | 38 | 12 | 15 | 5 | 6 | 17 | 91 |

B

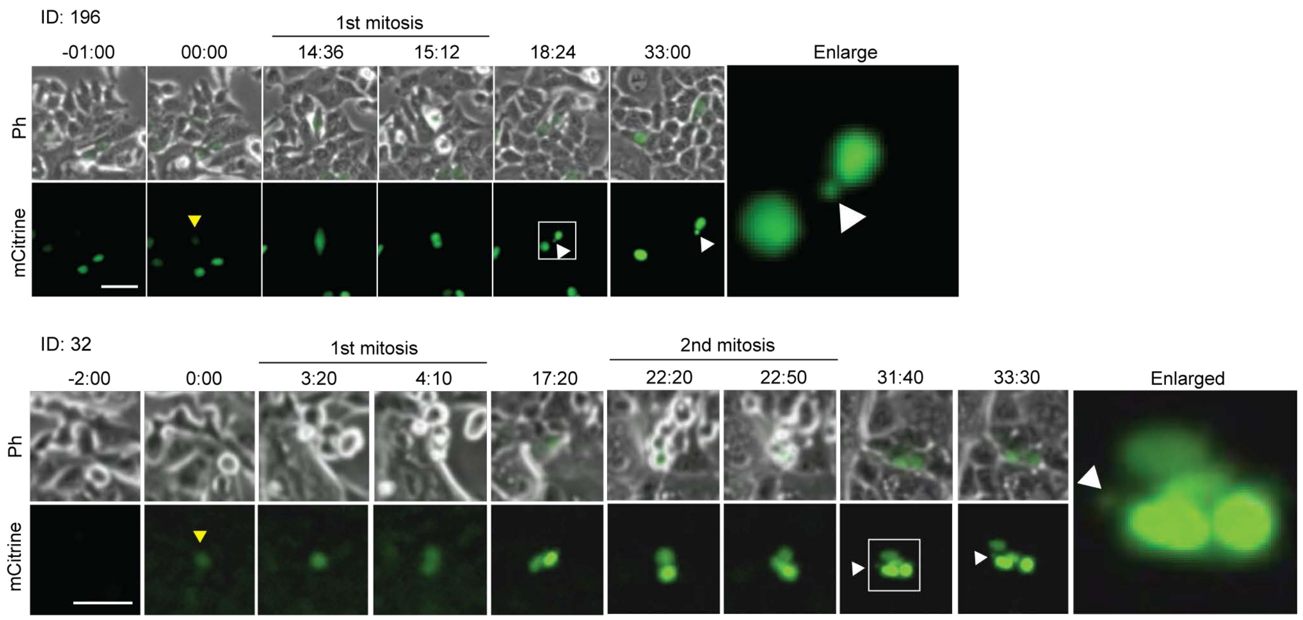


C

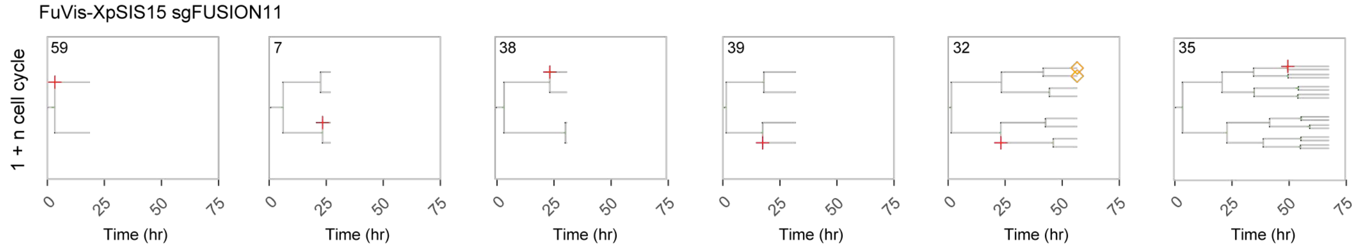


D

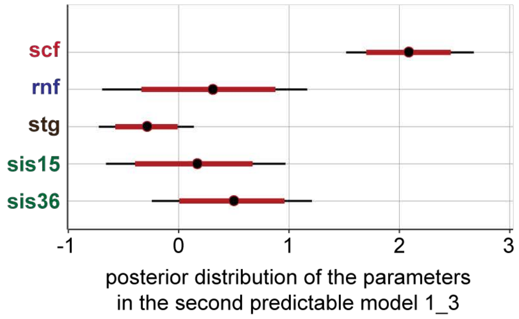


E

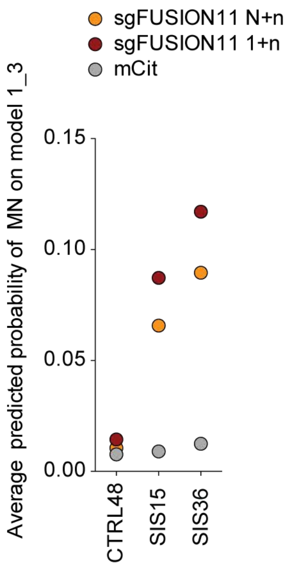

Supplementary figure 15 Kagaya et al.

A

$\mathbf{Int\_duration}_n \sim \text{LogNormal}(\mathbf{mu}_n, \mathbf{sigma}_j)$

| | | WAIC |
|---|---|---|
| Model 2_1 | $\mathbf{mu}_n = \mathbf{micro}*\text{MICRO}_n + \mathbf{scf}*\text{SCF}_n + \mathbf{rnf}*\text{RNF}_n + \mathbf{stg}*\text{Stage}_n + \mathbf{sis15}*\text{SIS15}_n + \mathbf{sis36}*\text{SIS36}_n + \mathbf{lin}_i$ | 17805.39 |
| Model 2_2 | $\mathbf{mu}_n = \mathbf{micro}*\text{MICRO}_n + \mathbf{scf}*\text{SCF}_n + \mathbf{rnf}*\text{RNF}_n + \mathbf{stg}*\text{Stage}_n + \mathbf{sis15}*\text{SIS15}_n + \mathbf{sis36}*\text{SIS36}_n + \mathbf{b}$ | 15754.86 |
| Model 2_3 | $\mathbf{mu}_n = \mathbf{scf}*\text{SCF}_n + \mathbf{rnf}*\text{RNF}_n + \mathbf{stg}*\text{Stage}_n + \mathbf{sis15}*\text{SIS15}_n + \mathbf{sis36}*\text{SIS36}_n + \mathbf{b}$ | 17934.47 |
| Model 2_4 | $\mathbf{mu}_n = \mathbf{micro}*\text{MICRO}_n + \mathbf{scf}*\text{SCF}_n + \mathbf{lin}_i$ | 18214.45 |
| Model 2_5 | $\mathbf{mu}_n = \mathbf{lin}_i$ | 18545.45 |
| Model 2_6 | $\mathbf{mu}_n = \mathbf{micro}*\text{MICRO}_n + \mathbf{scf}*\text{SCF}_n + \mathbf{b}$ | 16185.12 |

B

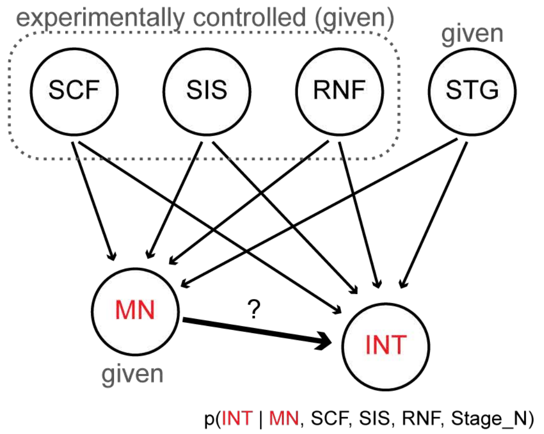


C

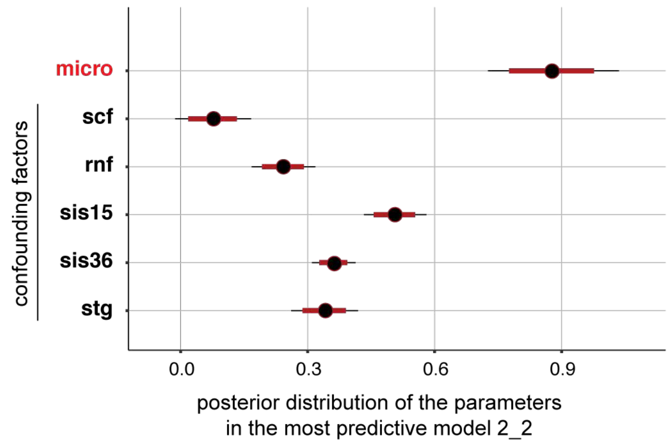
